# Integrative physiological, biochemical and transcriptomic analysis of hexaploid wheat roots and shoots provides new insights into the molecular regulatory network during Fe & Zn starvation

**DOI:** 10.1101/2021.04.03.438303

**Authors:** Om Prakash Gupta, Vanita Pandey, Ritu Saini, Tushar Khandale, Ajeet Singh, Vipin Kumar Malik, Sneh Narwal, Sewa Ram, Gyanendra Pratap Singh

## Abstract

In plants, iron (Fe) & zinc (Zn) uptake and transportation from the rhizosphere to the grain is a critical process regulated by complex transcriptional regulatory networks. However, understanding the combined effect of Fe & Zn starvation on their uptake and transportation and the molecular regulatory networks that control them lack in wheat. Here, we performed a comprehensive physiological, biochemical and transcriptome analysis in two bread wheat genotypes, *i.e.* Narmada 195 and PBW 502, differing in inherent Fe & Zn content to understand the mechanism of Fe & Zn homeostasis. Compared to PBW 502, Narmada 195 exhibited increased tolerance to Fe & Zn withdrawal by an increased level of antioxidant enzymes and DPPH radical scavenging activity along with less malondialdehyde (MDA), H_2_O_2_ level, increased PS accumulation and lower reduction of root and shoot Fe & Zn content and length, leaf chlorosis, and leaf area. By integrating physiological and biochemical data along with co-expression & functional genome annotation and gene expression analysis, we identified 25 core genes associated with four key pathways, *i.e.* Met cycle (10), PS biosynthesis (4), antioxidant (3) and transport system (8) that were significantly modulated by Fe & Zn withdrawal in both the genotypes. Genes of these four pathways were more considerably up-regulated in Narmada 195, allowing better tolerance to Fe & Zn withdrawal and efficient uptake and transportation of Fe & Zn. Chromosomal distribution and sub-genome wise mapping of these genes showed a contribution from all the chromosomes except group 5 chromosomes with the highest number of genes mapped to chromosome 4 (24%) and sub-genome D (40%). Besides, we also identified 26 miRNAs targeting 14 core genes across the four pathways. Together, our work provides a crucial angle for an in-depth understanding of regulatory cross-talk among physiological, biochemical and transcriptional reprogramming underlying Fe & Zn withdrawal in wheat. Core genes identified can serve as valuable resources for further functional research for genetic improvement of Fe & Zn content in wheat grain.

**Highlight:** Our work provides a crucial angle for a comprehensive understanding of the regulatory mechanism underlying Fe & Zn withdrawal associated with physiological, biochemical and transcriptional reprogramming in wheat.

## 1. Introduction

Micronutrients, especially iron (Fe) & zinc (Zn), although required in small quantities, play a pivotal role during the growth and development of plants and adversely affects the productivity of crops under deficient conditions (Rout and Sahoo, 2015). Fe & Zn deficiency in crop plants also results in a lower concentration of these nutrients in grains and leads to their deficiency in human beings after long-term consumption. Cognitive & immune impairment are the two significant Fe & Zn associated deficiency symptoms in humans that affect overall growth and development (Wishart, 2017). Globally, over two billion people are affected by Fe deficiency (Singh *et al.,* 2018), and around 30% population of developing countries are afflicted with Zn deficiency (Hotz and Brown, 2004). Fe deficiency-linked anaemia affects roughly 25% of the global population, leading to the loss of over 46,000 disability-adjusted life years (DALYs) in 2010 alone, and further, its deficiency caused mortality mainly in children under the age of 5 (Murray and Lopez, 2013). Therefore, researchers worldwide are ascertaining different methodologies to produce Fe & Zn enriched crops to ease the associated deficiency symptoms. Since wheat is the second most crucial cereal crop contributing considerably to food and nutritional security globally, consumption of Fe & Zn deficient wheat can lead to micronutrient deficiency in humans causing hidden hunger. Therefore, increasing the efficacy of Fe & Zn absorption, transportation and their accumulation in the wheat grain is required to enhance their grain content to mitigate their deficiency in human beings. This needs a better understanding of physiological, biochemical and molecular components associated with Fe & Zn metabolism in root and shoot compartments.

Plants adapt two prominent strategies for the uptake of Fe & Zn from the rhizosphere. Strategy I include direct uptake of Fe^2+^ and Zn^2+^ by ZRT-, IRT-like proteins (ZIPs) by enrichment of soil with protons (H^+^) and other reducing agents (Kobayashi and Nishizawa, 2012). Strategy II, generally used in graminaceous plants like wheat, operates *via* secretion of phytosiderophores (PSs), especially mugineic acid (MA), which chelate Fe^3+^ and the resulting Fe^3+^ - PS complexes are subsequently taken up by transporters like yellow stripe-like (YSL), ZIP *etc.* (Sperotto *et al.,* 2012; Connorton *et al.,* 2017a). Nicotianamine (NA), a metal chelator, mediates radial transport of Fe & Zn through the root to the shoot (Deinlein *et al.,* 2012). Several transporters including ZIP, YSL and metal tolerance protein (MTP) families have been predicted to facilitate the remobilization of Fe and Zn from leaves to the grains and from the maternal tissue into the endosperm cavity, aleurone and embryo (Tauris *et al.,* 2009) but their functional characterization is yet to be fully explained. Reports suggest considerable differences among wheat genotypes in tolerance to Fe & Zn deficiency (Hansen *et al.,* 1996; Cakmak *et al.,* 1994). Under both Fe & Zn deficiency, wheat plants exude higher PS content (Hansen *et al.,* 1996; Rengel *et al.,* 1998). Bread wheat synthesizes only one type of PS, *i.e.* 2’-deoxymugineic acid (DMA), where Met acts as a precursor (Mori and Nishizawa, 1987). DMA’s biosynthesis starts with the conversion of the methionine (Met) into SAM by s*-*adenosylMet (SAM) synthetase. Later, nicotianamine synthases (NAS) combine three SAM molecules to form one molecule of NA, which is then converted to an intermediate called 3′′-keto acid by NA aminotransferase (NAAT). This is followed by the synthesis of 2′-DMA by the subsequent action of a reductase, called DMA synthase (DMAS). Similarly, plants possess a complex antioxidant defence system comprising antioxidant enzymes and metabolites acting against reactive oxygen species (ROS) generated by several abiotic stresses, including metal deficiency (Kabir, 2016). Antioxidant enzymes, like superoxide dismutase (SOD), catalase (CAT), glutathione reductase (GR), peroxidase (POD), and ascorbate peroxidase (APX) *etc.*, are usually induced when plants are exposed to metal stress (Kabir, 2016). Therefore, understanding the role of antioxidative system genes in adaptation under Fe & Zn deficient conditions has become imperative. With the advancement of NGS technologies, several genome-wide approaches have been successfully exploited to study the global transcriptional changes under Fe and/or Zn stress in different crop plants, including maize (Mallikarjuna *et al.,* 2020), rice (Bandyopadhyay *et al.,* 2017; Zeng *et al.,* 2019) and wheat (Kaur *et al.,* 2019; Gupta *et al.,* 2020). Considerable variation in the expression profile of the genes mainly implicated in the Strategy II method of uptake was evident in crop plants when subjected to Fe and/or Zn deprivation (Quinet *et al.,* 2012; Bashir *et al.,* 2014; Kobayashi *et al.,* 2014; Li *et al.,* 2014). Similarly, Fe & Zn deficient condition in wheat plants have suggested the differential expression and essential role of many transcriptional regulators such as transcription factors (TFs) (Connorton *et al.,* 2017b; Sharma *et al.,* 2020) and critical gene(s) related to antioxidant enzymes, PS biosynthesis, and phytohormone homeostasis *etc.* (Schmidt *et al.,* 2000; Hindt and Guerinot, 2012). Despite the availability of advanced RNA sequencing (RNA-seq) based transcriptomic technologies and the availability of the sequenced wheat genome (IWGSC, 2014), very few reports are available to understand the molecular mechanism of wheat plant’s responses to Fe & Zn stress (Borrill *et al.,* 2018; Mishra *et al.,* 2019; Gupta *et al.,* 2020). Since Fe & Zn deficiency based appearance of phenotypes, especially their content in the grain, is the cumulative effect of physiological, biochemical and molecular reprogramming and studies deciphering the combined effect of these pathways in efficient and inefficient wheat genotypes under Fe & Zn deficiency is purely absent.

Therefore, in this investigation, two wheat genotypes, namely Narmada 195 and PBW 502 differing in grain Fe & Zn content, were used to understand the physiological, biochemical and molecular effect of Fe & Zn withdrawal in the nutrient solution. Twelve RNA-seq libraries generated from each genotype’s root and shoot tissues from different treatments represented RNAs expressed during control, partial (T1) and complete Fe & Zn withdrawal (T2). Besides analyzing the genome-wide expression profile of various transcripts associated with Fe-Zn metabolism, expression of 25 transcripts directly associated with PS biosynthesis, Met cycle, antioxidant system and the transport was also investigated using reverse transcription (RT)-qPCR to decipher the fold change variation during withdrawal. Overall, our results provide a unique and comprehensive insight into molecular, physiological and biochemical responses of two contrasting wheat genotypes under Fe & Zn withdrawal conditions.

## 2. Materials and Methods

### 2.1 Experimental setup-plant growth, Fe & Zn deficiency conditions and sampling

We selected two diverse hexaploid wheat varieties, namely Narmada 195 and PBW 502, in the current study as they exhibited different grain Fe & Zn content, with the former being more efficient than the latter (Table 1). The detailed outline of the experimental procedure is given in Fig. 1. Briefly, seeds were treated with 1% NaOCl for 10 min, followed by rinsing with distilled water three times. The sterilized seeds were subjected to cold stratification at 4^0^C overnight in the dark to break the dormancy, followed by germination for four days in Petri dishes containing three layers of moist and sterile Whatman filter paper. Subsequently, the seedlings were transferred to synthetic floater placed in polycarbonate Phyta Jar (75 x 74 x 138mm, Himedia, India) with 500 ml top size. We designed each synthetic floater with utmost care to accommodate nine seedlings at an equidistant position (Supplementary Fig. 1). Three treatments, *i.e.* control [C: Hoagland solution comprising full strength Fe (100 µM) & Zn (0.77 µM)], treatment 1 [T1: Hoagland solution comprising half-strength Fe (50 µM) & Zn (0.38 µM)] and treatment 2 [full strength for 18 days including four days germination on Petri plates followed by 0 µM Fe & Zn for the next 12 days], were used in the present study to see the differential response of Fe & Zn on contrasting wheat genotypes. Hoagland solution was prepared following the protocol of Yordem *et al.,* 2011 and frequently replaced on every alternate day. Seedlings were grown for 30 days in a growth chamber (E36H0; Percival Scientific Inc. Perry, IA) maintained with a 16 h day/8 h night cycle at 20±1 °C, 50–70% relative humidity, and a photon rate of μ300 mol quanta m^−2^s^−1^. Each genotype was represented with three independent biological replications for each treatment. For transcriptome, physiological and biochemical analysis, root and shoot tissues were independently harvested and snap-frozen in liquid nitrogen and stored at –80 °C until further use.

**Fig. 1:**
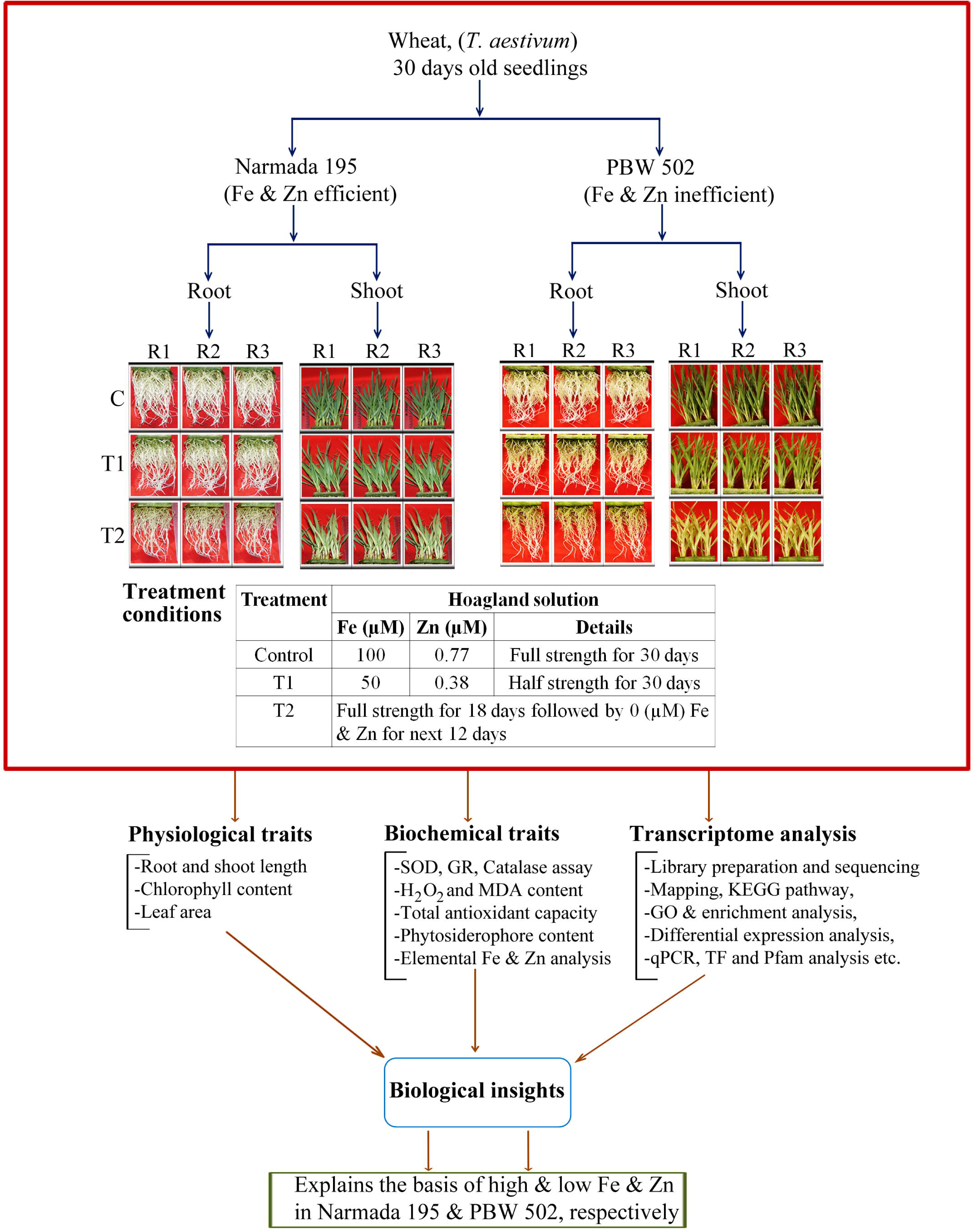
A schematic outline of the experimental setup, along with the analysed parameters.

**Table 1:** Pedigree details and Fe and Zn content of the genotypes used in the present study.

| Genotypes | Pedigree | Notification number and date | Average Grain Fe & Zn (ppm) | Special features | References |
| --- | --- | --- | --- | --- | --- |
| Narmada 195 | C306/HY65<br>(selection of N-112 ) | 19(E)<br>14.01.1982 | Fe:42;<br>Zn:38 | Tolerance to lodging | Gupta et al., 2018 |
| PBW 502 | W 485 /PBW 343//<br>RAJ 1482 | 161(E)<br>04.02.2004 | Fe:34;<br>Zn:30 | Resistance to yellow rust, brown rust and karnal bunt |  |

### 2.2 Physiological characterization

#### 2.2.1 Measurement of growth parameters

We harvested plants after 30 days of sowing grown under control, T1 and T2 conditions for measuring the different growth parameters. Root length was measured using Root Scanner (Model, Epson Perfection V 700 Photo, Win-RHIZO Programme V. 2009 c 32-bit Software) following the manufacturer’s protocol and shoot height was measured using a standard scale. After washing with deionized water, roots were placed on a 150 mm wide Petri plate filled with distilled water to manually observe the primary root and the first-order lateral root characteristics in each plant. To determine root characteristics, we used six wheat seedlings for each of the treatments, including three biological replicates. Each sample’s leaf area was determined using a Digital Leaf Area Meter (YMJ-C Series, China) following the manufacturer’s instructions. Three technical replicates were evaluated for each parameter.

#### 2.2.2 Determination of chlorophyll content and phytosiderophore assay

A chlorophyll meter (SPAD-502, Minolta, Japan) was used to estimate leaf chlorophyll concentration (SPAD value). Three biological replicates and three technical replicates per treatment were selected, and SPAD values were recorded from the fully matured leaves counted from the top of the plants.

Release of PS was estimated by Fe mobilization assay as per the method of Takagi, 1976. Briefly, nine seedlings from each of three biological replicates representing control, T1 and t2 conditions were undertaken for PS release assay under aeration for 4h in 20 ml deionized water. The assay was performed after 2h of exposure to a light period. After that, 8 ml of the collection solution was added to 0.5ml of 0.5M Na acetate buffer (pH 5.6), followed by 2 ml of the freshly prepared solution of Fe(OH)_3_. The resulting solution was stirred for 2h and filtered using Whatman #1, and the filtrate was mixed with 0.2ml of 6N HCl. The mixture was treated with 0.5ml of 8% hydroxylamine-hydrochloride by heating at 60°C for 20 min to reduce ferric iron. 1ml of 2M Na-acetate buffer (pH 4.7) and 0.2ml of 0.25% ferrozine was added into the above solution, and absorbance was taken at 562 nm as per the method of Khobra and Singh, 2018.

### 2.3 Biochemical characterization

#### 2.3.1 Enzymatic assays and determination of antioxidant capacity

Enzymes related to the antioxidant activity such as CAT, SOD and GR were estimated in seedlings at 30 DAS. Briefly, 200 mg seedlings were ground in liquid nitrogen and thoroughly homogenized in 1.2ml of 0.2M potassium phosphate buffer (pH 7.8 with 0.1mM EDTA) followed by centrifugation at 15,000×g for 20 min at 4°C. The supernatant was collected in fresh tube, and the pellet was redissolved in 0.8ml of the same buffer and centrifuged at 15,000×g for 15 min. The resulting supernatants were combined with the first extracts and stored at 4°C and used to estimate the different antioxidant enzyme activities.

The CAT activity was analyzed in a 3ml assay mixture comprising 2.94ml of 50mM phosphate buffer (pH 7.0), 50µl supernatant and 10µl of 30% H_2_O_2_. Once supernatant is added, the extinction coefficient of H_2_O_2_ (40mM^−1^cm^−1^ at 240 nm) was used to calculate the enzyme activity and expressed in terms of millimoles of H_2_O_2_ decomposed m^−1^ gFW^−1^ (Aebi and Lester, 1984). For measurement of the activity of SOD, 100 ul of supernatant was mixed in a 2.5ml reaction mixture comprising 50mM phosphate buffer (pH 7.8) (constituting 2mM EDTA, 9.9mM L-Met, 55μM NBT, and 0.025% Triton-X100) and 400μl of 1mM riboflavin. The reaction was started by illuminating the samples for 10 min, followed by recording the absorbance at 560nm instantaneously after the reaction was stopped. The enzyme activity (gFW^−1^) was estimated from a standard curve obtained using pure SOD (Beyer and Fridovich, 1987). For GR activity determination, 100µl of supernatant was mixed with 3 ml of assay mixture comprising 0.75mM DTNB, 0.1mM NADPH, and 1mM GSSG (oxidized glutathione). The reaction was initiated by the addition of GSSG, and the upsurge in absorbance (412nm) was recorded for 5 min. The GR activity was calculated using an extinction coefficient of TNB (14.15M^−1^cm^−1^) and expressed in millimole TNB m^−1^ gFW^−1^ (Smith *et al.,*1988).

For determination of hydrogen peroxide (H_2_O_2_), 500mg of tissue were homogenized with 5.0ml trichloroacetic acid (TCA) (0.1%, w/v, 4°C) and centrifuged at 12,000g for 15 min. The absorbance (390nm) was recorded by mixing 0.5ml of supernatant with 0.5ml of 10 mm potassium phosphate buffer (pH 7.0) and 1ml of 1M KI. The results were expressed as µmolg^−1^ FW using the extinction coefficient of 0.28μ cm (Velikova *et al.,* 2000). Malondialdehyde (MDA) content was estimated as previously described (Kosugi and Kikugawa, 1985). Briefly, wheat tissues were finely ground in 20% TCA with 0.5% 2-thiobarbituric acid (TBA) followed by heating at 95°C for 30 min and centrifugation for 10 min at 5000×g at 25°C. The supernatant was used to measure the absorbance at 532nm and 600nm, and MDA content was calculated by deducting the non-specific turbidity at 600nm by its molar extinction coefficient (Kosugi and Kikugawa, 1985). DPPH (2,2-diphenyl-1-picrylhydrazyl) free radical assay (Brand-Williams *et al.,* 1995) measured the percentage of antioxidant capacity (AA%) of 30 days old wheat tissues. The reaction mixture contained 0.5ml of the seedling sample, 3ml of absolute ethanol and 0.3ml of DPPH radical solution diluted to 0.5mM in ethanol. After 10 min. of incubation, the change in colour from deep violet to light yellow was recorded at 517nm and the DPPH scavenging activity was then calculated according to the formula given by Mensor *et al.,* (2001).

#### 2.3.2 Determination of Fe & Zn concentration

For estimating Fe & Zn content, root and shoot tissues were carefully washed with CaSO_4_ (1mM) and deionized water followed by drying in an oven at 80°C for two days (Kabir *et al.,* 2015). The dried tissues (0.5g) were mixed in 7ml HNO_3_ and digested in a micro-digestion system following the Organic B program (Microwave digestion system, Anton Paar). The concentration of Fe & Zn was then analyzed by Flame Atomic Absorption Spectroscopy (ECL, India). Standard solutions of Fe & Zn were separately prepared from their respective concentration of stock solutions.

### 2.4 Statistical analysis

Three independent biological replications for each sample were used for physiological and biochemical analysis. Data were subjected to one-way ANOVA by IBM SPSS package v. 20 (IBM Corp., New York, USA), and means were compared by Duncan’s multiple range test at a 5% significance level (P < .05).

### 2.5 RNASeq analysis

#### 2.5.1 RNA isolation, cDNA library construction, and Illumina sequencing

For transcriptome analysis, thirty days old seedlings (root and shoot) were used for RNA isolation and sequencing on the Illumina HiSeq4000 platform (Genotypic Technology Pvt. Ltd., Bangalore). RNA extractions were performed independently from each of the three biological replicates. Total RNA was extracted from root and shoot of all the three treatment conditions, *i.e.* C, T1 and T2, using the Qiagen RNeasy Plant Mini kit (Netherland) as per the manufacturer’s instruction. The yield and purity of RNA were evaluated by Nanodrop1000 spectrophotometer (NanoDrop, USA) and 1% denaturing RNA agarose gel, respectively. Before library preparation, the quality of all the RNA samples was thoroughly evaluated by Bioanalyzer 2100 (Agilent, USA) to ensure >8.5 RNA integrity number (RIN). RNA-seq library was prepared using NEBNext® Ultra™ directional RNA library prep kit (New England BioLabs, MA, USA) following the manufacturer’s instruction. The sequencing libraries were quantified by Qubit fluorometer (Thermo, USA) followed by an analysis of fragment size distribution on Agilent 2200 Tapestation (Agilent, USA). The average mean of the fragment size of all the libraries was 465 bp. The 2×150 bp chemistry was used for sequencing on the Illumina HiSeq4000 platform to produce on an average of 45.7 million raw sequencing reads per library.

#### 2.5.2 RNA-Seq data processing and assessment of differential gene expression

To ensure high-quality clean reads, we performed adopter trimming, removing low quality reads (QV <30 Phred score) and ambiguous N nucleotides (reads with unknown nucleotides ‘N’ >5%) using Trimmomatic-0.39 (Bolger *et al.,* 2014). The raw reads quality was ensured by FastQC (http://www.bioinformatics.babraham.ac.uk/projects/fastqc/). Finally, high-quality processed (QV>30), paired-end reads were de-novo assembled using a graph-based approach using rnaSPAdes program (Bankevich *et al.,* 2012). To reduce the redundancy without excluding sequence diversity required for further transcript annotation and the differential expression analysis, assembled transcripts were clustered using CD-HIT-EST (Fu *et al.,* 2012) with 95% similarity between the sequences. Processed reads from each library were mapped back to the final assembly using Bowtie2 (Langmead and Salzberg, 2012) with end-to-end parameters. DESeq R package (Anders and Huber, 2010) was used for differential expression analysis. Uneven sequencing library size and depth bias among the samples were scrubbed by library normalization using size factor calculation in DESeq. Log2 fold change (FC) values >1 were considered up-regulated, whereas those with an FC <1 were down-regulated. For their significant expression, these genes were further analyzed considering the statistical significance (*P*<0.05) and the false discovery rate (FDR 0.05) after Benjamin–Hochberg corrections for multiple testing. Clustv (http://biit.cs.ut.ee/clustvis/) was employed to construct heat maps for selected differentially expressed genes (DEGs) using the normalized expression values of genes.

#### 2.5.3 Gene Annotation, filtering and functional enrichment analysis of differentially expressed genes (DEGs) along with SSR mining

Gene annotations and functional enrichment analysis were carried out using multiple databases (GO term, Uniprot, KEGG pathway, Pfam, and PlnTF) to identify DEGs from both the root and shoot samples of C, T1 and T2 conditions and were significantly enriched in GO terms or biological pathways. Clustered transcripts were annotated using the homology approach to assign functional annotation using BLAST (Altschul *et al.,* 1990) tool against viridiplantae and *Triticum* proteins from UniProt database with e value <e^−5^ and minimum similarity >30%. Pathway analysis was performed using KAAS server (Finn *et al.,* 2015) by considering *Oryza sativa japonica* (Japanese rice), *Zea mays* (maize), *Musa acuminate* (wild Malaysian banana) and *Dendrobium catenatum* as reference organisms. Gene annotations against the GO database (http://geneontology.org/) were performed using the Blast2Go program. GO terms and biological pathways against the KEGG database (http://www.genome.jp/kegg) with a *p*-value < 0.05 were deemed to be significantly enriched in DEG analysis. Additionally, SSRs were identified in each transcript sequence using default parameters of MISA Perl script with simple repeat motif length ranging from monomer to hexamer (Finn *et al.,* 2016). To comprehend the conserved domains, Pfam Scan was used to predict the Pfam domain. Transcripts encoding TF were identified by homology search against known plant TFs from Plant TFdb. Venn diagram (VENNY 2.1) was constructed to highlight unique and common transcripts among genotypes, tissues, and treatments. Volcano Plot was created using Volcano Plot (https://paolo.shinyapps.io/ShinyVolcanoPlot/). An interaction network of selected genes was created using STRING 11.0.

#### 2.5.4 Quantitative reverse transcription PCR (RT-qPCR) validation

To further cross-check the reliability of RNA-Seq expression results, RT-qPCR was performed using the SYBR Green (Maxima SYBR Green qPCR Master Mix, Thermo Fisher Scientific) on a C1000^TM^ Touch Thermal Cycler (CFX96^TM^, BioRad). Briefly, total RNA isolated from all the treatment conditions, *i.e.* the C, T1 and T2, was treated with DNase I (Thermo Scientific) to remove DNA contamination in each sample. The first-strand cDNA was synthesized using the Maxima First-strand cDNA synthesis kit (Thermo Scientific) following the manufacturer’s guidelines. For qPCR, twenty-five differentially expressed genes belonging to Fe & Zn metabolism, including Met cycle, PS biosynthesis, antioxidant and transport system, were selected for experimental validation in response to various treatments of Fe & Zn withdrawal (Table S1). ADP-ribosylation factor 1 (ARF1) and actin were used as an internal control for expression normalization. As shown in Table S1, specific primers for real-time PCR were designed by Primer Premier 5.0 software (Premier Biosof International). All amplification programmes were: 95°C for 5min, followed by 40 cycles at 95°C (15s), 58°C (20s) with fluorescent signal recording at 72°C for 30s. Three independent biological replicates with three technical replicates were performed on each cDNA. The relative expression levels of genes were calculated using the formula 2^-ΔΔ^Ct^^ (Livak and Schmittgen, 2001). The student’s *t*-test (*P* < 0.05) was conducted to evaluate the significance of mean values.

#### 2.5.5 Identification of miRNAs and *in silico* functional characterization of Fe & Zn transport-related genes

Since miRNAs are critical regulators of gene expression during every stage and condition of the plant life cycle, we mined miRNAs from the selected transcripts obtained from RNA-Seq data regulating Fe & Zn metabolism, including Met cycle, PS biosynthesis, antioxidant system and transporters. The nucleotide sequences were manually retrieved from fastaseq file and used as a query sequence to mine the corresponding miRNA on psRNATarget: A Plant Small RNA Target Analysis Server (2017 Update) (https://plantgrn.noble.org/psRNATarget/home). The query sequences were used against published wheat miRNAs following the preset default programme. Further, transmembrane helix and topology analysis were performed to characterize the possible nature of essential protein using HMMTOP, MEMSAT3 and MEMSAT-SVM programs from the PSIPRED server. Also, the plausible subcellular localization of proteins was dogged using TargetP server (http://www.cbs.dtu.dk/) (Emanuelsson *et al.,* 2000), MEMSAT SVM and ProtComp v. 9.0. FFPred 3 program (PSIPRED server) was used to determine gene ontology domains *viz.* molecular function, biological process, and cellular component.

#### 2.5.6 Accessions numbers

The data generated from this study have been deposited in the NCBI Sequence Read Archive (SRA) database and are accessible with the submission ID-SUB6954440 and BioProjectID-PRJNA605691.

## 3. Results

### 3.1 Fe & Zn withdrawal agonies physiological growth and phytosiderophore release in wheat

Fe & Zn starved seedlings of wheat genotypes developed distinct phenotypic responses observed at 30 DAS. A significant decrease in the total leaf area was observed in both the genotypes under T1 and T2 conditions, but the decline was more prominent in inefficient genotype PBW 502 (51% and 49% for T1 and T2, respectively) compared to efficient genotype Narmada 195 (33% and 35% for T1 and T2, respectively) (Table 2). Compared to control, we observed a significant decrease in the shoot length and enhanced chlorosis in both the genotypes under T1 and T2 conditions, with more prominent symptoms in PBW 502 (Fig. 2A & B; Table 2). The reduction of shoot length was more significant in genotype PBW 502 where a decline of 44% and 51% was observed compared to 38% and 40% in Narmada 195 for T1 and T2, respectively (Table 2). The chlorophyll content decreased significantly in both the genotypes compared to controlled conditions, as is evident by the progressive chlorosis symptoms developed under T1 and T2. Compared to Narmada 195, we observed a steeper reduction of chlorophyll content in PBW 502, *i.e.* 38% and 81% under T1 and T2 conditions, respectively (Table 2). Compared to the control condition, there was a significant decrease in the number of lateral roots in both genotypes under T1 and T2 conditions (Fig. 2C & D). There was no substantial change in root length as measured in both the genotypes under the T1 condition. Moreover, the root length increased slightly in Narmada 195 and decreased in PBW 502 under the T2 condition compared to the control condition (Table 2).

**Fig. 2:**
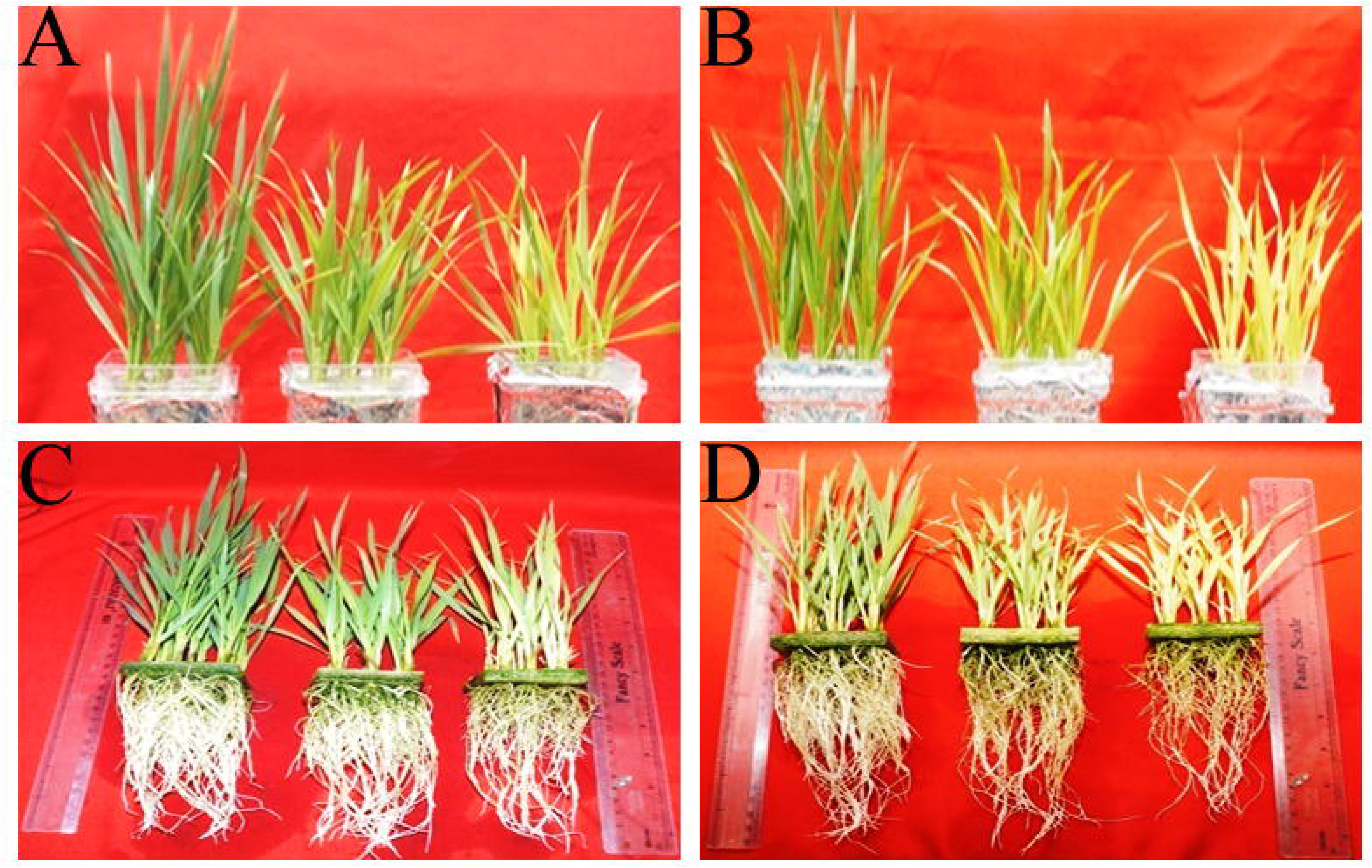
Figures representing the progression of morphological symptoms depicting chlorosis of leaves in (A) Narmada 195 and (B) PBW 502; and reduction in the number of lateral roots in (C) Narmada 195 and (D) PBW 502.

**Table 2:** Morpho-physiological features and Fe & Zn content (ppm on a dry weight basis) in Narmada 195 and PBW 502 grown under all three treatment conditions, *i.e.* C, T1 and T2.

| Treatments | Root Length<br>(cm) | Shoot length<br>(cm) | Chlorophyll<br>conc.<br>(SPAD value) | Leaf<br>Area<br>(cm<sup>2</sup>) | Zn (ppm) |  | Fe (ppm) |  |
| --- | --- | --- | --- | --- | --- | --- | --- | --- |
|  |  |  |  |  | Root | Shoot | Root | Shoot |
| Narmada 195 |  |  |  |  |  |  |  |  |
| Control (C) | 13.2±1.24<sup>ab</sup> | 25.9±0.21<sup>e</sup> | 40.2±2.41<sup>f</sup> | 9.3±0.58<sup>c</sup> | 29.3±5.24<sup>c</sup> | 19.4±4.83<sup>c</sup> | 228.3±13.26<sup>e</sup> | 192.4±28.70<sup>e</sup> |
| T1 | 14.5±0.91<sup>abc</sup><br>(9%) | 16±0.20<sup>c</sup><br>(38%) | 26.4±0.89<sup>e</sup><br>(34%) | 6.2±0.38<sup>b</sup><br>(33%) | 23.5±6.86<sup>abc</sup><br>(20%) | 13.5±2.54<sup>bc</sup><br>(30%) | 186.2±14.10<sup>d</sup><br>(18%) | 123.5±6.15<sup>d</sup><br>(36%) |
| T2 | 14.7±0.36<sup>bc</sup><br>(11%) | 15.7±0.33<sup>c</sup><br>(40%) | 15.3±0.53<sup>b</sup><br>(62%) | 6±0.21<sup>b</sup><br>(35%) | 22±5.23<sup>abc</sup><br>(25%) | 12.3±2.63<sup>ab</sup><br>(37%) | 79.2±10.03<sup>c</sup><br>(65%) | 58.8±5.76<sup>b</sup><br>(69%) |
| PBW 502 |  |  |  |  |  |  |  |  |
| Control (C) | 15.8±0.44<sup>c</sup> | 17.7±1.12<sup>d</sup> | 28.8±0.83<sup>d</sup> | 6±0.07<sup>b</sup> | 25.6±3.72<sup>bc</sup> | 15.4±2.11<sup>bc</sup> | 199±18.12<sup>d</sup> | 168.6±10.57<sup>e</sup> |
| T1 | 16±0.93<sup>c</sup><br>(1%) | 10±0.22<sup>b</sup><br>(44%) | 18±1.49<sup>c</sup><br>(38%) | 3.1±0.12<sup>a</sup><br>(51%) | 18.2±4.50<sup>ab</sup><br>(29%) | 10±3.86<sup>ab</sup><br>(34%) | 132.2±17.57<sup>b</sup><br>(34%) | 92.6±7.16<sup>c</sup><br>(46%) |
| T2 | 12.8±1.31<sup>a</sup><br>(19%) | 8.7±0.45<sup>a</sup><br>(51%) | 5.5±0.73<sup>a</sup><br>(81%) | 2.9±0.23<sup>a</sup><br>(49%) | 14.5±4.68<sup>a</sup><br>(43%) | 6.1±3.21<sup>a</sup><br>(60%) | 50±11.56<sup>a</sup><br>(76%) | 31.1±5.12<sup>a</sup><br>(82%) |

Note: Different letters indicate significant differences between means ± SD of treatments (n = 3) at P<0.05. Figures in brackets depict percentage change from respective control in each genotype.

On the other hand, estimation of Fe & Zn content in roots and shoots of both the genotypes suggested a significant decrease under both T1 and T2 conditions (Table 2); however, the decline was more evident in PBW 502 as compared to Narmada 195. The inefficient genotype PBW 502 suffered more reduction in Fe concentration in both roots and shoots under T1 (34%, 46% respectively) and T2 (76%, 82% respectively) (Table 2). Zn content decreased more in shoot than root, indicating a severe effect on Zn’s translocation in shoot under stress. Zn deficiency caused a decline of 34% in T1 and 60% in T2 of PBW 502 compared to controlled conditions (Table 2). Moreover, PS quantification showed that roots of Narmada 195 released more PS than roots of PBW 502 when plants were subjected to nutrient-deficient conditions, *i.e.* T1 and T2, with a maximum release under T2 condition (Table 3). This kind of favoured PS release in Narmada 195, even under T2 condition, signifies the genotypes dependent responses to efficient Fe and Zn transport and remobilization over other less efficient wheat genotypes.

**Table 3:** Changes in SOD, CAT, GR, H_2_O_2_, MDA and PS content and antioxidant capacity in Narmada 195 and PBW 502 in response to all the three treatment conditions, *i.e.* C, T1 and T2.

| Treatments | SOD (units/g FW) | CAT (μmoles/min/g FW) | GR (mM TNB min/g FW) | H<sub>2</sub>O<sub>2</sub> (μmol/g FW) | MDA (μM/ g FW) | DPPH Radical Scavenging activity (%) | PS (nmol of Fe equivalent /g root biomass) |
| --- | --- | --- | --- | --- | --- | --- | --- |
| **Narmada 195** |  |  |  |  |  |  |  |
| **Control** | 23.1±1.92<sup>d</sup> | 8.6±0.91<sup>a</sup> | 1.8±0.28<sup>ab</sup> | 2.4±0.27<sup>a</sup> | 4.7±0.77<sup>a</sup> | 30.7±1.82<sup>a</sup> | 2.7±0.19<sup>a</sup> |
| **T1** | 19.4±0.97<sup>c</sup><br>(-1.19) | 14.2±1.17<sup>b</sup><br>(1.65) | 4.4±0.88<sup>c</sup><br>(2.4) | 4.7±0.39<sup>b</sup><br>(1.95) | 5.9±0.24<sup>b</sup><br>(1.25) | 51.1±1.42<sup>d</sup><br>(1.66) | 3.6±0.82<sup>b</sup><br>(1.32) |
| **T2** | 7.7±2.32<sup>b</sup><br>(-3) | 18.6±0.91<sup>c</sup><br>(2.61) | 4.8±1.06<sup>c</sup><br>(2.6) | 6.4±0.49<sup>c</sup><br>(2.66) | 5.3±0.47<sup>ab</sup><br>(1.12) | 53.6±1.53<sup>d</sup><br>(1.74) | 6±0.22<sup>c</sup><br>(2.21) |
| **PBW 502** |  |  |  |  |  |  |  |
| **Control** | 21.3±0.94<sup>cd</sup> | 10.1±1.68<sup>a</sup> | 1.2±0.036<sup>a</sup> | 2.3±0.49<sup>a</sup> | 5±0.25<sup>ab</sup> | 37.4±1.41<sup>b</sup> | 2.1±0.21<sup>a</sup> |
| **T1** | 9.8±1.05<sup>b</sup><br>(-2.17) | 10.5±2.14<sup>a</sup><br>(1.03) | 2.4±0.35<sup>ab</sup><br>(2.0) | 3.9±0.29<sup>b</sup><br>(1.69) | 7.4±0.56<sup>c</sup><br>(1.48) | 40.3±1.59<sup>b</sup><br>(1.08) | 2.7±0.18<sup>a</sup><br>(1.32) |
| **T2** | 2.8±0.36<sup>a</sup><br>(-7.6) | 10.9±0.34<sup>a</sup><br>(1.07) | 2.7±0.59<sup>b</sup><br>(2.25) | 4.1±0.67<sup>b</sup><br>(1.78) | 8.1±0.89<sup>c</sup><br>(1.62) | 44.4±4.34<sup>c</sup><br>(1.19) | 3.6±0.30<sup>b</sup><br>(1.72) |

Note: Different letters indicate significant differences between means ± SD of treatments (n = 3) at P<0.05. The figure in brackets depicts fold change from respective control in each genotype.

### 3.2 Reactive oxygen species (ROS) and antioxidant scavenging system triggered during Fe & Zn withdrawal in wheat root and shoot

Reactive oxygen species and enzymes (SOD, CAT and GR) related to the antioxidant system were measured in seedlings to evaluate the effect of Fe & Zn withdrawal on their activity in both the wheat genotypes grown under C, T1 and T2 conditions. The result showed that the activity of SOD decreased significantly in both Narmada 195 and PBW 502 under both T1 (1.19-fold in Narmada 195; 2.17-fold in PBW 502) and T2 (3-fold in Narmada 195; 7.6-fold in PBW 502) conditions (Table 3). At the same time, the activity of CAT increased significantly in Narmada 195 under T1 (1.65-fold), and T2 (2.16-fold) compared to PBW 502 genotype where the increase (1.03-fold in T1 and 1.07-fold in T2) was not significant (Table 3). In contrast, GR and total antioxidant activities (DPPH radical scavenging activity) increased in both Narmada 195 and PBW 502 under Fe & Zn deficiency in both treatments, but the increase was more noticeable in Narmada 195 (Table 3). Content of H_2_O_2_ and MDA increased significantly in both the genotypes with a more pronounced effect in Narmada 195 (Table 3).

### 3.3 Transcriptome analysis of root and shoot of contrasting wheat genotypes in response to Fe & Zn withdrawal

To dissect the molecular cross-talk of Fe & Zn uptake and remobilization in root and shoot, we performed RNA-Seq analysis of two contrasting wheat genotypes (Narmada 195: efficient; PBW 502: inefficient) differing in total grain Fe & Zn content. Transcriptome data analysis resulted in an average of 43 million, ranging from 39 to 54 million, quality-filtered reads with an average ≥Q30 score of 94.52% (Supplementary Table S2). Around 77.5% of filtered reads were aligned back to the clustered transcripts (Supplementary Table S2). *De novo* assembly generated 176125 transcripts with a total length of 145224130 bp and an average length of 824 bp (Supplementary Table S3). The maximum transcripts were in size range of 300-500 bp (46.5%), followed by 1k-5k bp (23.7%) and 500-800 bp (21.6%) (Supplementary Table S3). Differential expression analysis was performed using the DESeq R package (Anders and Huber, 2010). Using a threshold value of logFC>1 for up-regulation, <1 for down-regulation with FDR of <0.05, T2 condition showed higher number of up (45518 & 25780 in root and 50735 & 17687 in shoot in Narmada 195 and PBW 502, respectively) and down (64205 & 22221 in root and 56618 & 19547 in shoot in Narmada 195 and PBW 502, respectively) regulated transcripts in both the tissues in Narmada 195 compared to T1 condition (up-regulated: 22462 & 48521 in root and 16617 & 53355 in shoot; down-regulated: 18646 &57414 in root and 19406 & 55597 in shoot in Narmada 195 & PBW 502, respectively) (Fig. 3A). Compared to control, there was exclusive induction of a significant number of transcripts in both the genotypes under T1 and T2 conditions, with the maximum in Narmada 195 (Fig. 3A). Next, we observed that 121 (C), 124150 (T1) and 75433 (T2) genes were shared in both root and shoot of both the wheat genotypes along with greater tissue and genotype-specific expression patterns (Fig. 3 B-D). Compared to PBW 502, Narmada 195 showed the larger number of tissue-specific expression pattern with the maximum in root under both T1 (2471: root; 872: shoot) (Fig. 3C) and T2 (7845: root; 6639-shoot) conditions (Fig. 3D; Supplementary Fig. S2 A&B). We observed 155242 shared transcripts in both the genotypes across the tissues and treatment condition (Supplementary Fig. S2C). Interestingly, this core set of shared DEGs showed significant differences in actual expression level in both the genotypes across the tissues and treatments (Fig. 3E-H; Supplementary Fig. S3). The analysis of gene expression profile across all the tissues, treatments and genotypes showed a positive correlation among commonly expressed genes (R^2^ >0.65)(Fig. 3I).

**Fig. 3:**
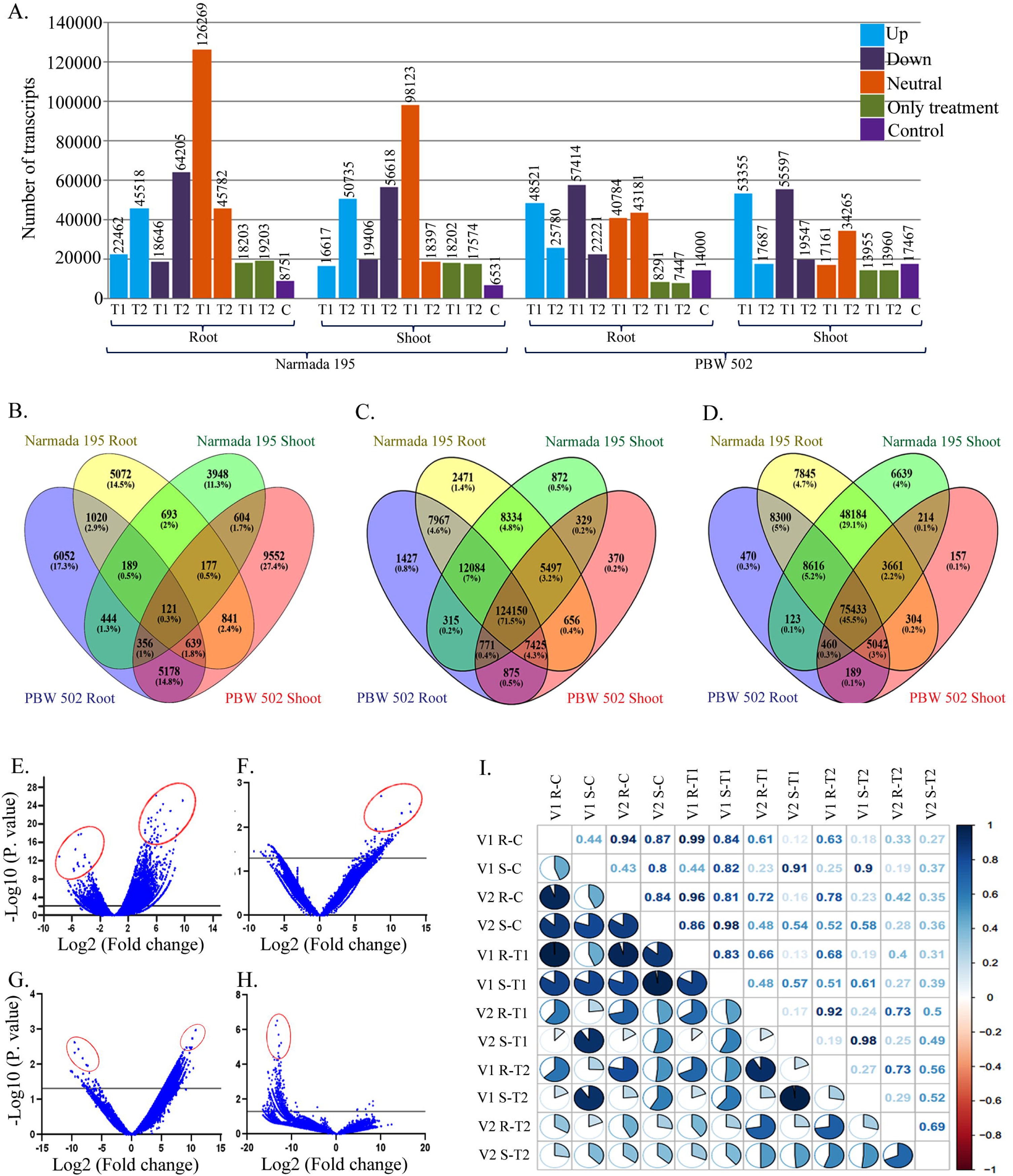
Analysis of DEG data from the wheat root and shoot during Fe & Zn withdrawal in Narmada 195 and PBW 502. (A) Comparison of the number of up (logFC>1), down (logFC< –1) and neutrally regulated (logFC= −1 to 1) DEGs under T1 and T2 conditions. (B) Venn diagram showing the numbers of unique and overlapping expressed genes in control, (C) T1, and (D) T2 across tissues and genotypes. (E) Volcano plot of DEGs in Narmada 195 T1 root & shoot; (F) Narmada 195 T2 root & shoot; (G) PBW 502 T1 root & shoot; (H) PBW 502 T2 root & shoot. Genes having the most significant differences are circled with red colour. (I) Pearson’s correlation matrix between samples using the cor R package.

### 3.4 Identification, functional classification, and enrichment analysis of DEG in response to Fe & Zn withdrawal in root and shoot

The GO annotation, classification and enrichment analysis of DEGs were performed to gain more insight into their potential involvement during biological, molecular, and cellular functions. Significant GO categories were assigned to all the DEG under both the treatment conditions, *i.e.* T1 and T2. Around 59.25% of the DEGs were functionally annotated against the Uniprot viridiplantae database. GO analysis revealed maximum categories in biological processes (1958 terms) followed by molecular function (1627 terms) and cellular component (523 terms) (supplementary table S4). The top ten over-represented significant terms of each of the three categories are given in Fig. 4A. ATP binding (11613 DEGs) and metal ion binding (3987 DEGs) were the most enriched GO term in the molecular function category, while integral components of the membrane (19337 DEGs) and nucleus (5768 DEGs) in the cellular component category and regulation of transcription (2192 DEGs) and translation (1942 DEGs) in biological process category (Fig. 4A). The transcripts’ E-value distribution showed that 58.77% of aligned transcripts had substantial similarity with an E-value <1e-60, whereas the remaining of the homologous sequences ranged from 1e-5 to 0 (Supplementary Fig. S4A). The similarity distribution in the reference showed that 47.45% of the sequences had a similarity higher than 80% (Supplementary Fig. S4B). Furthermore, we performed the Kyoto Encyclopedia of Genes and Genomes (KEGG) pathway (Xie *et al.,* 2011) of DEGs to identify critical pathways affected during T1 and T2 condition in both the wheat genotypes. From 80716 annotated transcripts in the KAAS server, we identified 206 pathways related to the plants’ various biological functions (Supplementary Table S5). Membrane trafficking (Ko04131) was the most abundant pathway in terms of the number of homologous transcripts, followed by chromosome and associated proteins (Ko03036) and exosome (Ko04147) (Fig. 4B). The maximum number of annotated DEGs were represented from *T. aestivum* (48%), followed by *T. Obliquus* (12%) and *N. nucifera* (7%) (Fig.4C; Supplementary Table S6).

**Fig. 4:**
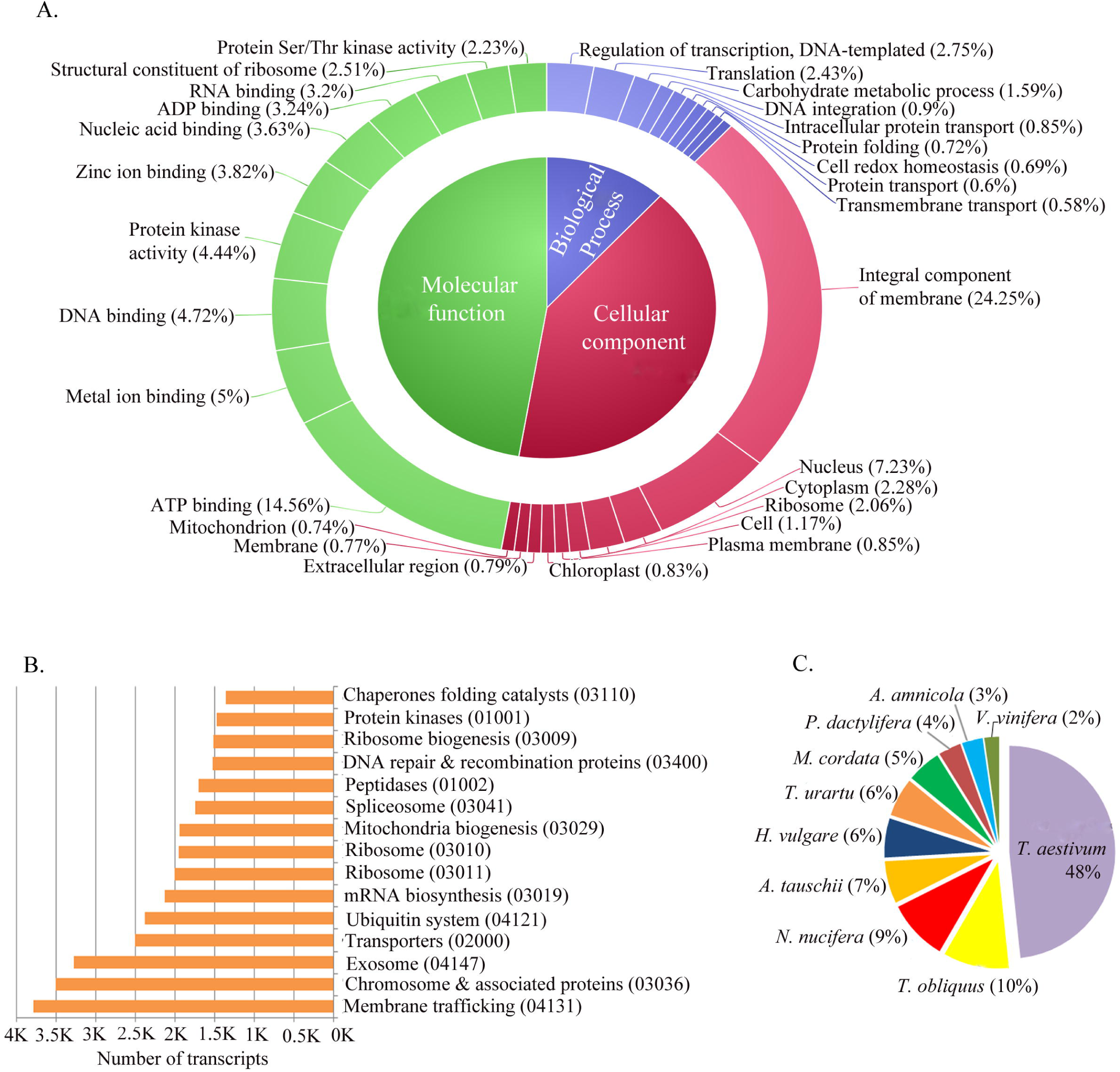
Gene Ontology (GO) categorization and enrichment analysis of DEGs.(A) Doughnut chart describing the frequency of top ten enriched GO terms under biological, cellular and molecular function. (B) Distribution of top 15 highly enriched KEGG pathways under T1 & T2 conditions. The *x*-axis shows the number of genes belonging to each KEGG pathway; the *y*-axis shows the pathways’ name. (C) A pie chart depicting the top ten species distribution for the annotated DEGs.

### 3.5 Identification and enrichment of DEGs associated with key Fe & Zn metabolic pathways including Met cycle, PS biosynthesis, antioxidant pathway and transport system

SAM, a substrate of the Met cycle, is used for the biosynthesis of PS that largely determine its accumulation and transport of Fe & Zn (strategy-II) throughout the plant system (Kobayashi and Nishizawa, 2012). Similarly, Fe & Zn stress also leads to the antioxidant pathway’s modulation (Cakmak *et al.,* 1997). Therefore, to uncover the key genes involved in these Fe & Zn related pathways, we performed the enrichment analysis of DEGs based on the KEGG pathway and identified 25 core genes (Table 4). Further, 10, 8, 4 and 3 genes were enriched in the Met cycle, Fe & Zn transport, PS biosynthesis and antioxidant pathway, respectively (Table 4). Next, we analyzed the chromosomal distribution of these genes in the wheat genome. Interestingly, except for group 5 chromosomes, all other chromosomes contributed to these genes, with the highest number of genes mapped to group 4 (24%) and group 7 chromosomes (20%). Group 2, 3 & 6 chromosomes contributed equally (16%) while chromosome 1 contributed only 4% (Fig. 5A). Further data analysis allowed us to map these core genes on different sub-genomes of wheat. The maximum number of genes were mapped on sub-genome D (40%) followed by sub-genome B (36%) and sub-genome A (24%) (Fig. 5B). As the interacting behaviour of the genes of linked pathways is crucial for proper substrate channelling, we performed STRING analysis to get more insights into the interacting nature and co-expression of the core genes. Excitingly, we observed significant (p-value <1.0e-16) protein-protein interaction (PPI) among themselves, signifying that these core genes have expressively more interaction among themselves than what would be expected from a random set of proteins of similar size, drawn from the genome (Fig. 5C). Such enrichment shows that the proteins are at least partially biologically connected as a group. We also dissected these core genes at the protein level by *in-silico* analysis of transmembrane helices, MW, Pi, GRAVY and protein types. The results showed a range of these parameters, with eight protein being membrane-bound on different cell organelles signifying their organelle-specific role during Fe & Zn homeostasis (Supplementary Table S7).

**Fig. 5:**
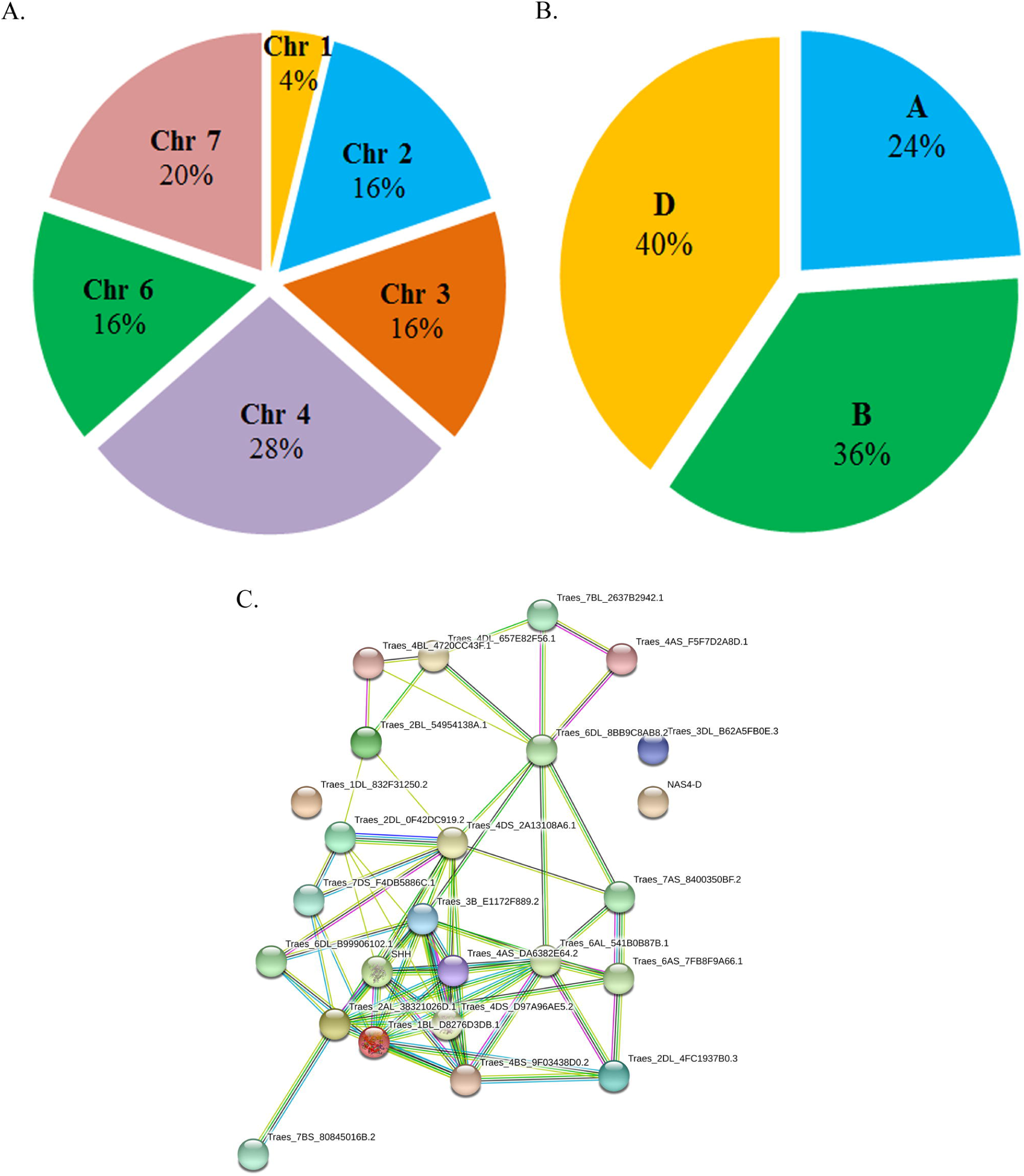
Genomic distribution of core genes. (A) Chromosomal distribution of the core DEGs. (B) Pie charts showing the subgenomic distribution of the core genes. A, B, and D refers to the sub-genome of hexaploid wheat. (C) STRING analysis of the co-expressed core DEGs showing protein-protein interaction (PPI) network.

**Table 4:** List of critical genes associated with Met cycle, PS biosynthesis, antioxidant pathway and Fe & Zn transport system and their chromosomal locations

| Sr. No | Transcript ID | IWGSC Gene Id | Chr location | Gene Name |
| --- | --- | --- | --- | --- |
| **Methionine cycle** |  |  |  |  |
| 1. | NODE_139907_length_361_cov_3.34314 | TraesCS4B02G014700 | 4B | Methionine synthase |
| 2. | NODE_10030_length_2203_cov_31.2784 | TraesCS2D02G493500 | 2D | S-adenosylhomocysteine hydrolase |
| 3. | NODE_34963_length_1154_cov_15.2402 | TraesCS4A02G065100 | 4A | Homocystein S-methyltransferase |
| 4. | NODE_16088_length_1799_cov_24.8939 | TraesCS3B02G228500 | 3B | S-adenosylmethioninesynthetase |
| 5. | NODE_3245_length_3201_cov_10.5372 | TraesCS6D02G202500 | 6D | S-adenosylmethioninedecarboxylas |
| 6. | NODE_88157_length_530_cov_3.56 | TraesCS2A02G396400 | 2A | 1-aminocyclopropane-1-carboxylic acid synthase (ACS2) |
| 7. | NODE_123078_length_398_cov_84.0525 | TraesCS7D02G029100 | 7D | 5'-methylthioadenosine nucleosidase |
| 8. | NODE_93542_length_503_cov_57.471 | TraesCS2D02G545000 | 2D | 5-methylthioribose kinase |
| 9. | NODE_17367_length_1739_cov_60.076 | TraesCS4D02G104900 | 4D | Methylthioribose-1-phosphate isomerase |
| 10. | NODE_119513_length_407_cov_38.3949 | TraesCS4B02G157700 | 4B | Cystathionine gamma-synthase |
| **Phytosiderophore biosynthesis** |  |  |  |  |
| 11. | NODE_63800_length_708_cov_24.6263 | TraesCS3B02G068500 | 3B | Nicotianamine synthase (NAS) |
| 12. | NODE_13301_length_1959_cov_96.1266 | TraesCS1B02G300600 | 1B | Nicotianamine aminotransferase(NAAT) |
| 13. | NODE_37066_length_1105_cov_18.341 | TraesCS2D02G313700 | 2D | DMAS or 3"-deamino-3"-oxonicotianamine reductase |
| 14. | NODE_23989_length_1468_cov_38.2831 | TraesCS7A02G159200 | 7A | 2'-deoxymugineic-acid 2'-dioxygenase or DMAD |
| **Fe and Zn transport system** |  |  |  |  |
| 15. | NODE_31063_length_1254_cov_6.90575 | TraesCS6D02G406400 | 6D | Solute carrier family 30 (zinc transporter) |
| 16. | NODE_30981_length_1256_cov_40.597 | TraesCS4A02G025400 | 4A | Solute carrier family 39 (zinc transporter) |
| 17. | NODE_23970_length_1469_cov_50.3607 | TraesCS7B02G251700 | 7B | ZIP metal ion transporter(ZTP29) gene |
| 18. | NODE_14782_length_1872_cov_6.75564 | TraesCS4D02G299400 | 4D | Natural resistance-associated macrophage protein 2 |
| 19. | NODE_32210_length_1224_cov_7.50043 | TraesCS4B02G190300 | 4B | Solute carrier family 25 (mitochondrial iron transporter) |
| 20. | NODE_35382_length_1144_cov_6.6685 | TraesCS7D02G413000 | 7D | Vacuolar iron transporter family protein |
| 21. | NODE_117812_length_412_cov_7.27171 | TraesCS1D02G295000 | 3B | Iron transport multicopper oxidase |
| 22. | NODE_137313_length_366_cov_1.34084 | TraesCS3D02G257900 | 3D | Multidrug resistance protein 1 homolog |
| **Antioxidant system** |  |  |  |  |
| 23. | NODE_15181_length_1849_cov_11.136 | TraesCS6B02G056800 | 6B | Catalase (CAT) |
| 24. | NODE_22558_length_1521_cov_15.6965 | TraesCS7A02G090400 | 7A | Super oxide dismutase (SOD) |
| 25. | NODE_19134_length_1658_cov_101.022 | TraesCS6A02G383800 | 6A | Glutathione reductase (GR) |

### 3.6 Differential expression analysis of core genes using RT-qPCR and RNA-Seq data associated with Fe & Zn homeostasis

Based on treatment and tissue conditions, we planned four different groups *viz.* (control Narmada 195 root *vs* treated Narmada 195 root; control Narmada 195 shoot *vs* treated Narmada 195 shoot and control PBW502 root *vs* treated PBW502 root; control PBW502 shoot *vs* treated PBW502 shoot) for DEG comparison. A heat map of the top 25 up and down-regulated DEGs from all the four groups is given in Supplementary Fig. S5a-d). Additionally, in response to Fe & Zn withdrawal, the expression landscape of up, down and neutrally regulated DEG in all the four groups is given in Supplementary Table S8-11.To confirm RNA-seq data’s reliability, we also performed a more rigorous expression measure for 25 selected genes using RT–qPCR analysis. We observed a good agreement with a high linear correlation (R^2^ >0.8; see supplementary Fig. S6) between RNA-seq and RT–qPCR technologies, suggesting RNA-seq analyses’ reliability. Interestingly, the expression of the enriched 25 genes associated with four pathways, *i.e.* Met cycle, PS biosynthesis, Fe & Zn transport, and antioxidant pathway, represented substantial differences in expression profile across the treatment and tissue conditions in both the genotypes (Fig. 6A & B; 7A & B). The expression pattern of enriched genes was highly induced in T2 compared to T1 in both the genotypes, but the level of gene stimulation was more predominant in the root of efficient genotype Narmada 195 compared to root and shoot of non-efficient PBW 502 (Fig. 6A & B; 7A & B; Supplementary Fig. S7). Further, heat map analysis of 25 core genes showed the formation of two clusters, each specific for T1 and T2 conditions across the tissues and genotypes (Supplementary Fig. S7). The expression pattern of all the homoeologues of four pathways’ core genes showed varying accumulation across the tissues and treatments (Supplementary Table S12). Amongst genes of the Met cycle, Met synthase (TraesCS4B02G014700) was found to be highly induced followed by methylthioribose-1-phosphate isomerase (TraesCS4D02G104900) and 5-methylthioribose kinase (TraesCS2D02G545000), suggesting their crucial role during flux channelling towards PS biosynthesis (Fig. 6A). The NAS (TraesCS3B02G068500) and NAAT (TraesCS1B02G300600) were among the highly up-regulated genes of the PS biosynthesis pathway, followed by DMAD (TraesCS7A02G159200) and DMAS (TraesCS2D02G313700) (Fig. 6B), which could be associated with efficient utilization of SAM substrate in Narmada 195 root. Amongst Fe & Zn transporter genes, solute carrier family 30 (zinc transporter) (TraesCS6D02G406400), Natural resistance-associated macrophage proteins *2* (NRAMP2) (TraesCS4D02G299400), multidrug resistance protein (MDRP) 1 homolog (TraesCS3D02G257900) and vacuolar iron transporter (VIT) family protein (TraesCS7D02G413000) were highly induced in root of Narmada 195 under T2 condition (Fig. 7A). The expression of GR (TraesCS6A02G383800) was highly induced, followed by SOD (TraesCS7A02G090400) and CAT (TraesCS6B02G056800) during Fe & Zn withdrawal conditions which could be associated with the triggering of the ROS pathway (Fig. 7B).

**Fig. 6:**
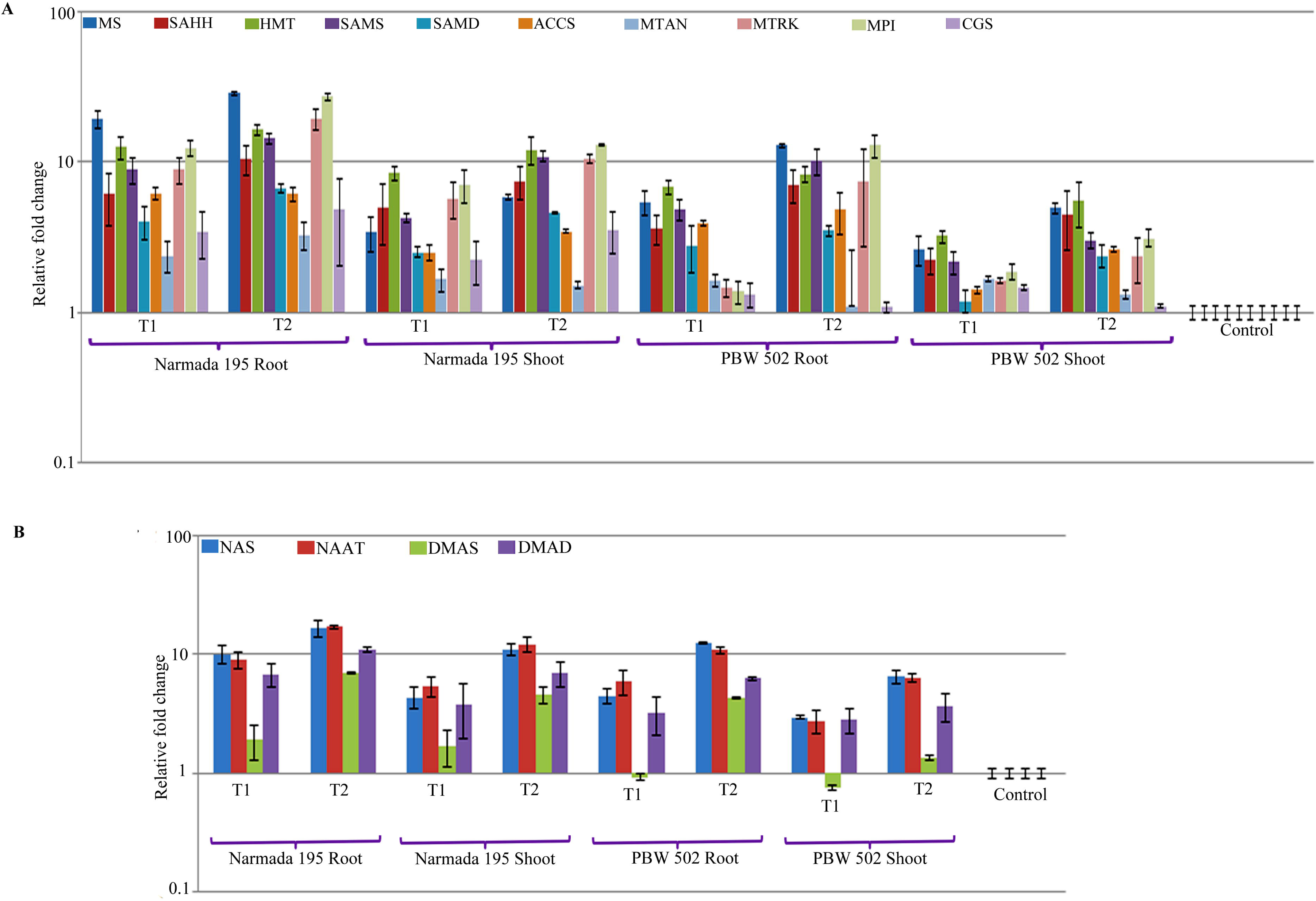
RT-qPCR validation of the core genes associated with (A) Met cycle and (B) PS biosynthesis. ARF1 and actin were used as an endogenous control for normalizing the Ct value. Data are means of three independent biological replicates (P ≤ 0.05, n = 3). Error bars represent the means ± SD (n = 3). Abbreviations: MS: Met synthase; SAHH: S-adenosylhomocysteine hydrolase; HMT: homocystein S-methyltransferase; SAMS: S-adenosylMetsynthetase; SAMD: S-adenosylMet decarboxylase; ACCS: 1-aminocyclopropane-1-carboxylic acid synthase (ACS2); MTAN: 5’-methylthioadenosine nucleosidase; MTRK: 5-methylthioribose kinase; MPI: methylthioribose-1-phosphate isomerase; CGS: cystathionine gamma-synthase; NAS: Nicotianamine synthase; NAAT:niconianamine amino transferase; DMAS: 3’’-deamino-3’’-oxonicotianamine reductase; DMAD: 2’-deoxymugineic-acid 2’-dioxygenase.

**Fig. 7:**
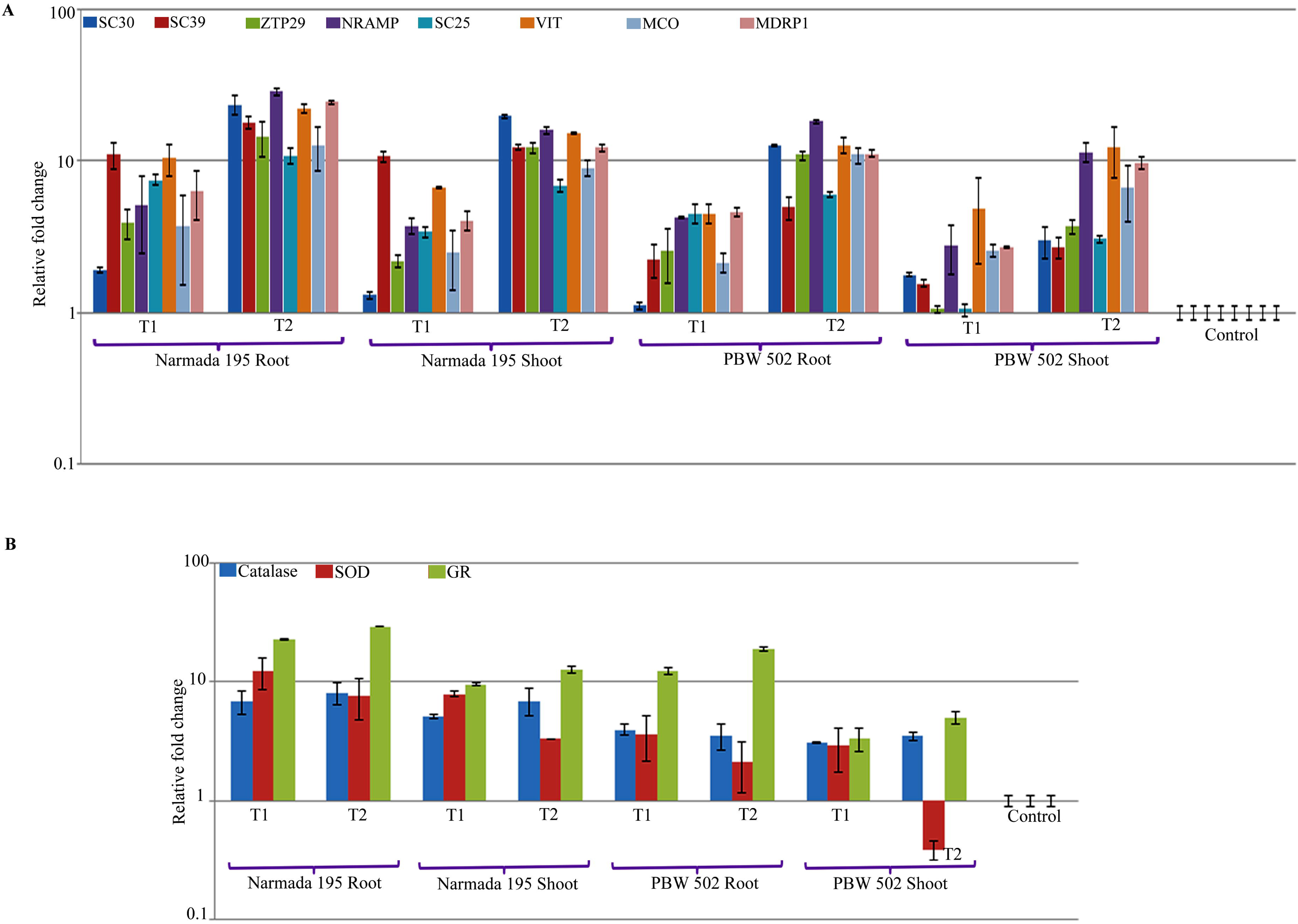
RT-qPCR validation of the core genes associated with (A) Fe & Zn transport system and (B) antioxidant pathway. ARF1 and actin were used as an endogenous control for normalizing the Ct value. Data are means of three independent biological replicates (P ≤ 0.05, n = 3). Error bars represent the means ± SD (n = 3). Abbreviations: SC30:solute carrier family 30 (zinc transporter); SC39: solute carrier family 39 (zinc transporter); ZTP29: ZIP metal ion transporter; NRAMP: natural resistance-associated macrophage protein 2; SC25: solute carrier family 25 (mitochondrial iron transporter); VIT: vacuolar iron transporter; MCO: iron transport multicopper oxidase; MDRP1: multidrug resistance protein 1 homolog; SOD: superoxide dismutase; GR: glutathione reductase.

### 3.7 Identification of transcriptional regulatory genes and protein families (Pfam) during Fe & Zn withdrawal

Several reports have evidenced the association of different transcription factors (TF) families including NAC, bHLH, EIN, PYE, MYB and WRKY during Fe & Zn homeostasis in *Arabidopsis* and rice model plants (Kobayashi *et al.,* 2007; Ogo *et al.,* 2007; Long *et al.,* 2010; Kobayashi and Nishizawa, 2012; Zamioudis *et al.,* 2014; Yan *et al.,* 2016; Wang *et al.,* 2019). To dissect the complex transcriptional regulatory network of Fe & Zn uptake, transport and remobilization, we identified vital genes encoding various TFs under T1 and T2 conditions. Based on RNA-Seq data, we identified 58 TF families (FDR ≤0.05) across the tissues and treatment conditions (Supplementary Table S13). Interestingly, further enrichment analysis of DEGs exhibited that bHLH (∼3461) was the most abundant TF family followed by NAC (∼2443), MYB (∼2203) and WRKY (∼2127), signifying their potential role during Fe & Zn homeostasis under both T1 and T2 conditions (Supplementary Table S13). Comparatively, Narmada 195 showed the highest number of up and down-regulated DEGs coding for different TF families (Fig. 8) under T2 condition, whereas PBW 502 exhibited the highest number of up- and down-regulated DEGs under T1 condition (Fig. 8). This demonstrated that Narmada 195 has efficient transcriptional reprogramming of the gene networks related to efficient uptake and transportation even under Fe and Zn withdrawal conditions, thereby increasing grain accumulation. We also performed protein family (Pfam) domain prediction and identified ∼4057 families across tissue and treatment conditions with the highest number of transcripts represented by PPR repeat (2913) followed by PPR repeat family (2025) and protein kinase domain (1702) (Supplementary Table S14).

**Fig. 8:**
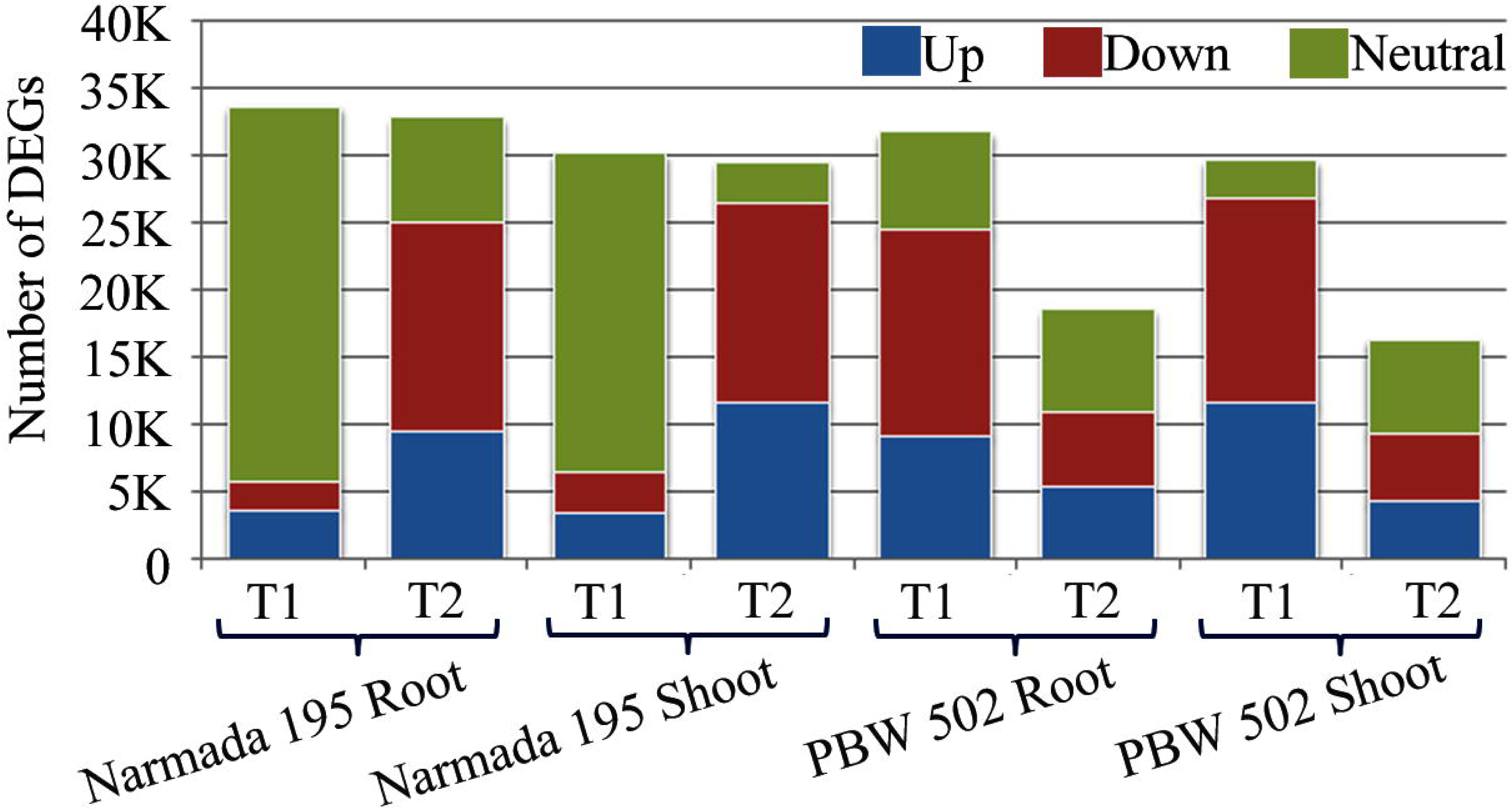
List of TFs significantly associated with Fe & Zn withdrawal (FDR ≤0.05) in root and shoot of the contrasting wheat genotypes. Blue bars represent up-regulated, the red bar is down-regulated, and the green bar represents neutrally regulated.

### 3.8 Identification of regulatory miRNAs and SSR markers from DEGs related to Fe & Zn homeostasis

In plants, miRNAs have appeared as prime regulator of many biotic and abiotic stresses, including low micronutrient availability by modulating the expression of transporter genes associated with nutrient uptake and mobilization (Gupta *et al.,* 2014a & 2014b; Paul *et al.,* 2015; 2016; Gupta *et al.,* 2020). To decode miRNAs’ involvement across the tissue and treatment conditions in efficient and inefficient wheat genotypes, we identified miRNAs targeting core genes of the Met cycle, PS biosynthesis, transporters, and antioxidant pathway. Result revealed 26 miRNAs targeting 14 core genes across all four pathways, while 11 genes did not show any corresponding miRNAs (Supplementary Table S15). Interestingly, the highest number of miRNAs were represented by methylthioribose-1-phosphate isomerase gene (4) followed by s-adenosylMet decarboxylase, natural resistance-associated macrophage protein and catalase, each targeted by three miRNAs (Supplementary Table S15). Likewise, the use of SSR markers in QTL mapping to understand the genetic basis of Fe & Zn accumulation in grains has immense potential to devise new breeding strategies for increasing grain micronutrient content through marker-assisted selection (Krishnappa *et al.,* 2017). SSR analysis of RNA-Seq data in the present study led us to identify 41723 SSRs, out of which 7147 transcripts were represented by over 1 SSR (Supplementary Fig. S8A). Identified SSRs were predominated by mono (32%), tri (28%) and tetra (27%) nucleotide repeats (11649) (Supplementary Fig. S8A). Alike, motif prediction of these SSRs showed the abundance of T/A followed by CAG/GTC and CTG/GAC with least represented by TCA TCG/AGT AGC (Supplementary Fig. S8B).

## 4. Discussion

Understanding the physiological, biochemical and molecular mechanism regulating Fe & Zn uptake, transport and remobilization have great potential for improving Fe & Zn content in wheat grain. In this study, we first performed a study of the dynamic changes in different physiological parameters such as root & shoot length, chlorophyll content and leaf area of two hexaploid wheat genotypes having different grain Fe & Zn content exposed to 50% (T1) and complete withdrawal (T2) of Fe & Zn. Secondly, the biochemical parameters related to the antioxidant system, PS content and Fe & Zn content were determined in both shoot and root. Thirdly, we performed transcriptome analysis with interaction network and transcriptional module that led to identifying a core set of genes involved in Fe & Zn homeostasis. By integrating physiological and biochemical data along with co-expression & functional genome annotation and gene expression analysis, we identified four key pathways, *i.e.* Met cycle, PS biosynthesis, antioxidant and Fe & Zn transport system, significantly affected by Fe & Zn deficiency (Fig. 9). The results presented here provide a comprehensive understanding of the gene regulatory network of four critical pathways associated with Fe & Zn uptake, transport and remobilization in wheat.

**Fig. 9:**
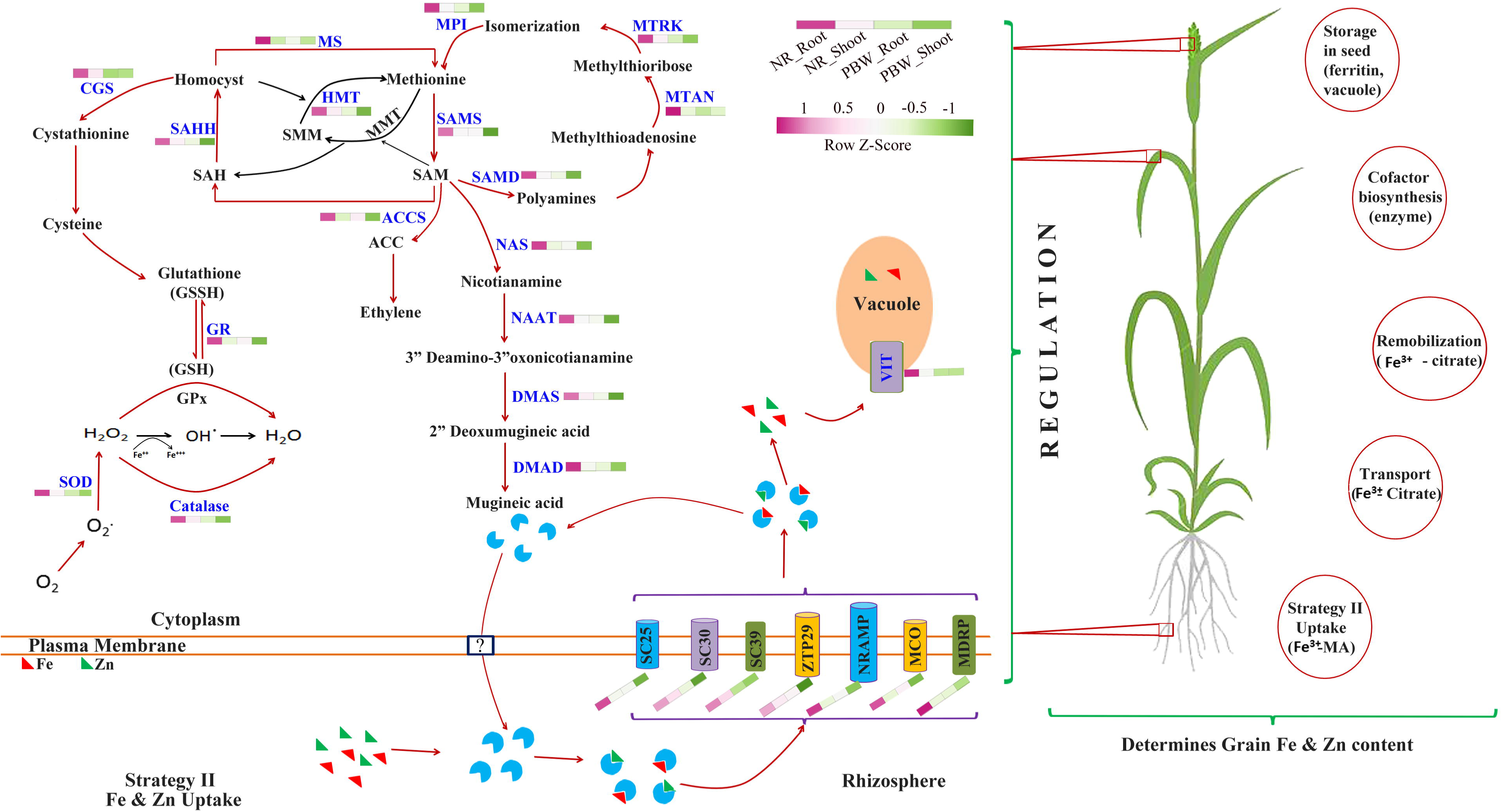
Schematic model representing the involvement of core genes in the Met cycle, PS biosynthesis, antioxidant and transport system in response to Fe & Zn withdrawal in wheat. The expression of transcripts is depicted in the colour scale. Colour scale refers to the log2 fold change values of differentially expressed transcripts: red colour refers to those transcripts positively regulated, while green colour denotes negatively regulated transcripts upon Fe & Zn withdrawal. NR: Narmada 195; PBW: PBW 502.

### 4.1 Fe & Zn withdrawal significantly modulates the physiological and anti-oxidant potential in wheat

Previous reports have shown the adverse effects of Fe & Zn starvation on physiological traits, including root and shoot growth (Garnica *et al.,* 2018). Here, the data demonstrated significant decreases in shoot growth under both T1 and T2 conditions. Leaf exhibited chlorosis after 14 days of growth in T1 and 6-7 days of treatment under T2 in both the genotypes compared to control. Iron deficiency based chlorosis of young leaves is a common symptom in plants (Santos *et al.,* 2019). Besides, micronutrients, especially Fe, are supposed to act through membrane stabilization by acting as a cofactor in several biological processes, including chlorophyll biosynthesis and photosynthesis (Ma *et al.,* 1999). The severity of physiological symptoms was more prominent in nutrient in efficient genotype PBW 502 than the efficient genotype Narmada 195. A significant decrease in the total leaf area was observed for both genotypes under T1 and T2, but the decline was more pronounced for PBW 502 (51% and 49% for T1 and T2, respectively) compared to Narmada 195 (33% and 35% for T1 and T2 respectively). Similarly, both Fe & Zn concentration decreased significantly in both shoot and root parts under nutrient deficiency stress (Table 2); however, the decline was more pronounced in PBW 502 than Narmada 195. Zn content decreased more in shoot than root, showing a severe effect on Zn’s translocation in shoot under stress. Similarly, Impa *et al.,* 2013 reported higher Zn content in the leaf of efficient rice genotype IR55179 than sensitive genotype under stress. Several root-related processes, such as efflux of PS, proton exudation and formation of Fe plaques, may influence higher Fe & Zn uptake in Fe & Zn efficient genotypes (Impa *et al.,* 2013; Rose *et al.,* 2013).

Activities of SOD, CAT and GR related to the antioxidant system were undertaken in the present study. There was a significant decrease in the activity of SOD in both Narmada 195 and PBW 502 under both T1 (1.19-fold in Narmada 195; 2.17-fold in PBW 502) and T2 (3-fold in Narmada 195; 7.6-fold in PBW 502) conditions. Based on metal cofactors in the active site, SODs are of three types, *i.e.* MnSOD, Cu/ZnSOD, and FeSOD. Fe and Zn are essentially required for the activity of Cu/ZnSOD and FeSOD. Deficiency in these two elements leads to reduced expression of Cu/ZnSOD and FeSOD isoforms. Literature suggests increased SOD activity in plants grown under Fe deficiency mainly because of enhanced expression of Cu/Zn or Mn-SOD isoform (Ranieri *et al.,* 2001; Molassiotis *et al.,* 2006; Donnini *et al.,* 2011). Several other reports also showed a decrease in the activity of SOD in wheat (Cakmak *et al.,* 1997), bean (Cakmak and Marschner, 1993) and cotton (Cakmak and Marschner, 1987). Interestingly, the decreases in activity of SOD because of Fe & Zn deficiency was prominent in inefficient genotype, as observed for PBW 502 in the current study. The variation in the amounts of physiologically active Fe & Zn present in the plants can be accounted for the differential severity of deficiency symptoms despite the nearly similar concentration of Fe & Zn in leaves. Possibly, efficient genotypes contain higher amounts of Fe & Zn chelators in tissues, such as S-containing amino acids, nicotianamine and PS, which influence the mobility of Fe & Zn in plants and enhance the physiologically active Fe & Zn pool at the cellular level (Stephan *et al.,* 1994; Graham and Welch,1996).

The increased CAT activity observed in efficient genotype Narmada 195 under Fe & Zn deficiency suggests that these ROS-scavenging antioxidant enzymes have a vital role in eliminating destructive oxidant species. Increased activity of CAT in response to Fe & Zn deficiency have also been reported in *Poncirus trifoliate* (Xiao *et al.,* 2010), Apple (Jin-Hua *et al.,* 2012), rice (Kabir *et al.,* 2017) and wheat (Sharma *et al.,* 2004). The notably increased GR activity observed in Narmada 195 suggested higher efficiency in converting O_2_ to H_2_O_2_ for protecting plants against oxidative stress. The Fe & Zn deficiency triggered enhanced GR activity might activate both antioxidant enzyme and ASC–GSH cycle, thus stimulating the synthesis of antioxidant metabolites. Differential expression analysis of SOD, CAT and GR during Fe & Zn withdrawal conditions exhibited a significant increase in their expression levels, which could be associated with the ROS pathway’s triggering (Fig. 7B). Increased expression of CAT and GR was correlated with higher activities of these enzymes. However, increased expression of the SOD gene could not lead to enhanced enzyme activity because Fe and Zn are required for the enzyme’s efficient functioning by acting as a cofactor.

Variations have also been reported for antioxidants associated with differential Fe & Zn efficiency in contrasting genotypes for other plants (Frei *et al.,* 2010; Santos, 2019). Compared to Narmada 195, we observed a substantial increase in MDA content in PBW 502 because of Fe/Zn withdrawal intolerance. An increase in MDA content shows the severity of stress experienced by any plant owing to its negative effect on the cell membranes (Chakraborty and Pradhan, 2012). The increase in MDA level in PBW 502 could be because of an overproduction of ROS *vis-à-vis* to an inadequate capacity to detoxify it. Scavenging of ROS for restoring redox metabolism, preserving cellular turgor, and structures actively function during abiotic stress in plants (Mittler, 2006). Therefore, inefficient wheat genotype PBW 502 could not activate Fe & Zn uptake mechanism as competently as efficient wheat genotype Narmada 195 and, therefore, suffers from more significant oxidative damage with a lower antioxidative response.

### 4.2 Fe & Zn withdrawal significantly modulates genes associated with the Met cycle

SAM synthesized from Met by SAM synthase serves as a precursor for the biosynthesis of polyamines, PSs and ethylene (Mori and Nishizawa, 1987). It is a metabolically very essential compound by acting as a methyl group donor in several biochemical reactions. Constant recycling of various metabolites of the Met salvage pathway is critical to maintaining the physiological level of Met. The Met cycle actively recycles Met to meet the augmented demand for PS biosynthesis (Ma *et al*. 1995). Amongst the enzymes associated with the Met cycle, formate dehydrogenase (FDH; Suzuki *et al.,* 1998), Fe-deficiency-induced protein 1 (IDI1; Yamaguchi *et al.,* 2000), and adenine phosphoribosyltransferase (Itai *et al*., 2000) are up-regulated because of Fe deficiency. Other Met cycle enzymes are also transcriptionally up-regulated under Fe & Zn deficient conditions (Kobayashi *et al.,* 2014; Gupta *et al.,* 2020). In our study, we identified ten genes associated with the Met salvage pathway (Fig. 9). Expression analysis showed up-regulation of almost all the genes in both the genotypes with the higher up-regulation in Fe & Zn efficient Narmada 195 root under T2 condition (Fig. 4A & 9). Increased up-regulation of most of the Met salvage pathway genes ensures a constant supply of various metabolites for the biosynthesis of PSs under Fe & Zn deficiency condition. These results imply that the Met salvage pathway’s activation might be one of the mechanisms for enhanced accumulation of PSs, leading to efficient uptake of Fe & Zn in efficient wheat genotype Narmada 195 even in Fe & Zn withdrawal conditions compared to inefficient PBW502 genotype. Furthermore, the Met salvage pathway genes have been shown to be contributed by different sub-genomes (Fig. 3A). This kind of sub-genomic distribution of the Met cycle’s essential genes suggests the importance of all genome under Fe & Zn withdrawal conditions. Based on these results, future work should be aimed at deciphering these genes’ molecular function using approaches like CRISPR/Cas9, TILLING or heterologous system to gain more insight into their role during Fe & Zn deficiency.

### 4.3 Phytosiderophore biosynthesis is negatively regulated by Fe & Zn withdrawal

Biosynthesis and release of PSs are very critical in strategy II mode of Fe & Zn uptake and translocation from rhizosphere to the grains (Fig. 9). Increased PS release from roots of graminaceous species exposed to micronutrient deficiency has been reported by several workers, including Fe deficiency (Khobra & Singh, 2019; Divte *et al.,* 2019) and Zn deficiency (Khobra & Singh, 2019; Ahmadzadeh and Khoshgoftarmanesh, 2019). However, there is limited information about the combined effect of Fe & Zn withdrawal in root and shoot comprehensively. The varying degree of PS accumulation in response to Fe & Zn deficiency is attributed to environmental conditions, physiological performance, genetic background and composition of nutrient medium (Arora *et al.,* 2019; Niyigaba *et al.,* 2019). In this investigation, efficient genotype Narmada 195 showed a higher accumulation of PS under T2 condition, *i.e.* complete Fe & Zn withdrawal, compared to inefficient PBW502 wheat genotype (Table 3). The differences in PS accumulation can be correlated with the differential expression of genes related to PS biosynthesis. In this investigation, the RT-qPCR expression analysis suggested up-regulation of genes related to PS biosyntheses such as NAS, NAAT, DMAS and DMAD, in both the genotypes, *i.e.* Narmada 195 and PBW502, under both treatment conditions, *i.e.* T1 and T2 (Fig. 4B & 9). However, compared to PBW 502, up-regulation of these genes was higher in Narmada 195 that might be one reason for more PS biosynthesis and release leading to efficient uptake and transportation of Fe & Zn. Various other reports have also shown that the PS biosynthesis genes are up-regulated under Fe and Zn (Ahmadzadeh and Khoshgoftarmanesh, 2019; Gupta *et al.,* 2020) deficiency conditions in different crops, including rice, wheat, barley and oats. Binding of Fe & Zn with PS shares similar biochemical confirmation and a similar regulatory mechanism of biosynthesis and/or release of PS under Zn & Fe deficiency (Rengel and Romheld, 2000). However, different divalent cations such as nickel have also been reported to compete with Fe & Zn in binding with available PS suggesting a series of regulatory cross-talk. Nickel deficiency significantly enhanced shoot Fe & Zn concentrations in wheat, while it decreased shoot Fe & Zn concentrations in triticale (Ahmadzadeh and Khoshgoftarmanesh, 2019). Therefore, it is imperative to revisit the binding kinetics of different divalent cations, including Fe & Zn, with the available PS to understand their uptake and transportation mechanism better. Higher accumulation of PS in Narmada 195 shows better Fe & Zn uptake and transportation even under T2 condition. Varietal differences in PS accumulation *vis-à-vis* grain Fe & Zn content can be utilized to decipher the molecular mechanism of Fe & Zn accumulation.

### 4.4 Transporters play a pivotal role to cope up with the Fe & Zn deficiency

Metal ion transporters play a crucial role in modulating different metal ions’ cellular homeostasis, including Fe & Zn. To maintain the precise metallic homeostasis in plants, several gene families of metal ion transport such as cation diffusion facilitator (CDF) family, NRAMP, MDRP, VIT, ZIP, and P-type ATPase take part in the uptake and transport of metal ions by plants (Zhang *et al.,* 2013; Connorton *et al.,* 2017a). SLC30 (ZnTs) and SLC39 (ZIP) transporters (Fig. 9), identified in our study, have been widely reported as two major Zn transporters of the CDF family regulating the cellular Zn homeostasis in mammals (Cotrim *et al.,* 2019). Both the transporters regulate cellular Zn homeostasis by trafficking the Zn in the opposite direction, *i.e.* SLC30 mediates Zn efflux out of the cytosol into the extracellular space or intracellular compartments while SLC39 imports Zn into the cytosol from extracellular space or intracellular compartments (Kimura and Kambe, 2016). While SLC39 is ubiquitously expressed in mammalian tissue (Gaither and Eide, 2001), the expression of SLC30 vary greatly across tissue and developmental condition (Schweigel-Röntgen, 2014). We, for the first time, identified SLC30 and SLC39 associated with Zn homeostasis in wheat. Differential expression analysis also exhibited higher expression of these two transporters under T2 condition in both the tissues, *i.e.* root and shoot in Narmada 195 compared to PBW502, showing their possible association with efficient mobilization of Zn in Narmada 195. Our *in-silico* analysis of SLC30 and SLC39 showed six transmembrane helices (Supplementary table S7), a typical feature of these proteins (Huang and Tepaamorndech, 2013). Further investigations are required in other model plant system for comprehensive functional characterization, including their possible competition/interaction between Fe & Zn. We also identified an SLC25 transporter localized at the mitochondrial membrane for iron transport (Supplementary table S7). On the other hand, ZTP29, another ZIP initially identified on endoplasmic reticulum (ER) in *Arabidopsis* under salt stress (Wang *et al.,* 2010), is homologous to *Ta*ZIP16 in wheat (Evens *et al.,* 2017). Zn deficiency induces the expression of *Ta*ZIP16 in wheat (Evens *et al.,* 2017), which supports our results of enhanced expression of ZTP29 in both the genotypes with higher expression in Narmada195 genotype under both Fe & Zn withdrawal condition. Members of ZIPs have emerged as a critical membrane transporter family in Zn’s journey from soil to seed (Palmgren *et al.,* 2008). Zn deficiency-induced expression of *Ta*ZIP16 and ZTP29 might be involved in balancing the Zn status of mitochondria even under Zn withdrawal condition. Further work on cross-talk of ZTP29 expression in response to both Fe & Zn in different tissues and developmental condition could enhance our present understanding of Fe & Zn metabolism in plants.

NRAMP family is another widely characterized transport protein ranging from bacteria to human for their crucial role during transport of various divalent cations, including Fe & Zn. (Garrick *et al.,* 2006). Different members of the NRAMP family have shown differential expression behaviour in response to Fe & Zn deficiency conditions in various crops, including rice (Peris-Peris *et al.,* 2017), sorghum and maize (Wairich *et al.,* 2019), wheat (Gupta *et al.,* 2020), suggesting their crucial role in maintaining cellular Fe & Zn homeostasis. We found a higher expression of NRAMP transporter in Narmada 195 in both root and shoot compared to PBW 502 (Fig. 9). Despite these initial results, comprehensive work on genome-wide identification and characterization of all the NRAMP family members would enhance our current understanding of its role during the transport of Fe, Zn and other divalent cations. As a part of a safe Fe storage strategy, vacuolar sequestration is another essential mechanism that regulates Fe homeostasis. VITs play a central role in sustaining the optimal physiological range of Fe (Aggarwal *et al.,* 2018). The expression pattern of VIT1 and VIT2 differed in rice and wheat in response to varying level of Fe & Zn (Sharma *et al.,* 2020). *Os*VIT1 remained unaltered to post seven days of Fe starvation, while *Os*VIT2 showed significant down-regulation (Zhang *et al.,* 2012) in rice. Hexaploid wheat genome has two *VIT* genes, *i.e. TaVIT1* and *TaVIT2*, and have differential expression pattern across the tissues (Connorton *et al.,* 2017b). Recently, Sharma *et al*. (2020) have shown up-regulation of vacuolar-iron transporters like (VTL) genes in wheat under Zn, Mn and Cu deficiency. In contrast, our result showed up-regulation of *VIT* in root and shoot of both the genotypes under T1 and T2 condition. This type of differences in results could be attributed to the difference in species, genotype, treatment condition and duration of treatment, age of plants, *etc*.

### 4.5 Multiple transcription factors are involved in Fe & Zn uptake and transportation in wheat

Over the past 20 years, significant efforts were made in identifying numerous transcriptional regulators of Fe & Zn in plants (Connorton *et al.,* 2017a). Several TF families, including bHLH NAC, C2H2, MYB, WRKY and bZIP (Assuncao *et al.,* 2010; Kim *et al.,* 2012), have widely been characterized for their crucial role in nutrient uptake and homeostasis in model plants such as *Arabidopsis* and rice. In the recent past, the transcriptomic approach has emerged as one of the critical methods to identify and characterize the TF families in wheat exposed to either Fe (Kaur *et al.,* 2019; Wang *et al.,* 2019), Zn (Wang *et al.,* 2017) or Fe & Zn together (Mishra *et al.,* 2019; Gupta *et al.,* 2020). Taken together, our result also showed that bHLH, NAC, MYB, WRKY were among the top TF families showing induced expression in response to Fe & Zn withdrawal in both the wheat genotypes (Fig. 6; Supplementary Table S12). Similarly, bZIP was also one of the highly induced TF families detected in the current study. Higher accumulation of two significant Zn deficiency-induced TF, such as bZIP19 and bZIP23, was associated with up-regulation of the essential PS biosynthesis gene NAS and ZIP members transporter (Clemens *et al.,* 2013; Inaba *et al.,* 2015). Despite significant progress in understanding the molecular network and cross-talk during Fe & Zn homeostasis, future research should be directed towards using the reverse genetic and functional complementation test in the model organism.

### 4.6 Post-transcriptional regulation plays a critical role in regulating core genes of Fe & Zn homeostasis

Post-transcriptional regulators such as miRNA play a central role in regulating the gene expression associated with various biotic, abiotic and nutrient homeostasis in plants (Gupta *et al.,* 2014a; 2014b; Paul *et al.,* 2015). During the last decade, several reports have witnessed miRNAs’ involvement in maintaining the cellular Fe & Zn homeostasis in various plants (Paul *et al.,* 2016; Zeng *et al.,* 2019; Gupta *et al.,* 2020). However, reports on Fe & Zn deficiency’s combined effect on miRNA cross-talk, especially in wheat, is limited (Gupta *et al.,* 2020). In the current study, we identified 26 miRNAs targeting four critical pathways, *i.e.* Met cycle, PS biosynthesis, transport and antioxidant system (Supplementary Table S15). As in our previous report (Gupta *et al.,* 2020), these miRNAs might play a critical role in regulating the transcript abundance of the cellular Fe & Zn homeostasis pathway, which could be decisive in controlling the level of grain Fe & Zn in wheat. Further work on Spatio-temporal expression profiling of miRNAs and their corresponding target genes and their functional validation in response to Fe & Zn withdrawal needs to be performed either in wheat or any model plants. This will help develop the miRNA-based gene regulatory network to better understand the molecular mechanisms of Fe & Zn homeostasis at the level of post-transcription.

## 5. Conclusion

Combined analysis of physiological, biochemical and molecular parameters in two wheat genotypes, *i.e.* Narmada 195 and PBW 502, under both T1 and T2 conditions showed the adaptive superiority of Narmada 195 over PBW 502 for efficient uptake and mobilization of Fe & Zn from rhizosphere to grains. An increase in the antioxidant capacity and less severity of Fe & Zn deficiency symptoms in Narmada 195 signifies that high antioxidant response plays a crucial role in Fe-Zn deficiency tolerance in Narmada 195. Moreover, PBW 502 could suffer more because of an overproduction of ROS and, at the same time, an inadequate capacity to detoxify it. Higher expression of genes related to the Met cycle, PS biosynthesis, antioxidant and transport system in Narmada 195 indicates better Fe & Zn uptake and transportation even under T2 condition (Fig. 9). The study has contributed significantly to our current understanding of physiological, biochemical and molecular mechanisms of Fe & Zn uptake and translocation under the varying level of Fe & Zn in wheat genotypes. This will help design strategies to improve Fe and Zn content in wheat under deficient conditions to boost grain nutritional quality through Fe & Zn bio-fortification programmes.

## Supplementary data

**Fig S1:** Flowchart giving an outline of the hydroponic experiment.

**Fig. S2:**Venn diagram showing the numbers of unique and overlapping differentially expressed genes in root and shoot.

**Fig. S3:**Volcano plots showing the expression levels of the genes

**Fig. S4:** (A) Pie chart showing the e-value distribution plot (B) Graph showing the percentage similarity distribution.

**Fig. S5a:** Heat map showing the expression pattern of top-up and down-regulated genes in efficient Narmada 195 root.

**Fig.S5b:** Heat map showing the expression pattern of top-up and down-regulated genes in efficient Narmada 195 shoot.

**Fig. S5c:** Heat map showing the expression pattern of top-up and down-regulated genes in inefficient PBW502 root.

**Fig.S5d:** Heat map showing the expression pattern of top-up and down-regulated genes in inefficient PBW502 shoot.

**Fig. S6:** Scatter plot represents fold changes in gene expressions measured by RNA-seq and an RT-qPCR assay of core genes.

**Fig. S7:** Heat map of core genes associated with four key pathways.

**Fig. S8:** Identification of (A) SSRs and their (B) motif prediction in transcriptomic data.

**Table S1:** List of primers used for qPCR in the present study.

**Table S2:** Summary of read statistics of RNA-Seq libraries.

**Table S3:** Assembly statistics of RNA-Seq libraries.

**Table S4:** Gene Ontology (GO) analysis of DEGs in response to Fe & Zn withdrawal condition

**Table S5:** List of enriched KEGG pathway identified using KAAS server.

**Table S6:** Statistics of species distribution of annotated DEGs.

**Table S7:** *In-silico* analysis of core genes for various parameters at the protein level.

**Table S8:** List of DEGs in response to Fe & Zn withdrawal in the root of efficient wheat genotype Narmada 195.

**Table S9:** List of DEGs in response to Fe & Zn withdrawal in the shoot of efficient wheat genotype Narmada 195.

**Table S10:** List of DEGs in response to Fe & Zn withdrawal in the root of inefficient wheat genotype PBW502.

**Table S11:** List of DEGs in response to Fe & Zn withdrawal in the shoot of inefficient wheat genotype PBW502.

**Table S12:** Expression profiles of genes/gene families involved in the Met cycle, PS biosynthesis, Fe & Zn transport system and antioxidant pathways.

**Table S13:** List of TF families enriched across tissues, genotypes and treatment conditions.

**Table S14:** List of protein families (Pfam) enriched across tissues, genotypes and treatment conditions.

**Table S15:** Identification of the corresponding miRNA of core genes during Fe & Zn homeostasis.

## Acknowledgements

Authors are thankful to the Indian Council of Agricultural Research, Department of Agricultural Research and Education, Govt. of India for providing financial help under grant no. 1006422, institutional project (CRSCIIWBRSIL 201500900190) and Department of Biotechnology, Govt. of India under the grant BT/NABI-Flagship/2018. We also acknowledge Sendhil R and Vijay Singh’s technical help in statistical analysis and growing the seedlings, respectively.

## Author contributions

OPG, VP and SR conceived the program, designed the experiment; RS, TK, AS, VKM, OPG, VP and SN performed the experiments; OPG analyzed the transcriptome data, OPG, RS, VP and AS analysed physiological and biochemical data; OPG prepared the figures; OPG and VP wrote the manuscript; SR and GP supervised the writing; All authors read and approved the final manuscript.

